# Resting brain fluctuations are intrinsically coupled to visual response dynamics

**DOI:** 10.1101/650291

**Authors:** Michaël E. Belloy, Jacob Billings, Anzar Abbas, Amrit Kashyap, Wen-ju Pan, Rukun Hinz, Verdi Vanreusel, Johan Van Audekerke, Annemie Van der Linden, Shella D. Keilholz, Marleen Verhoye, Georgios A. Keliris

## Abstract

How do intrinsic brain dynamics interact with processing of external sensory stimuli? We sought new insights using functional (f)MRI to track spatiotemporal activity patterns at the whole brain level in lightly anesthetized mice, during both resting conditions and visual stimulation trials. Our results provide evidence that quasiperiodic patterns (QPPs) govern mouse resting brain dynamics. QPPs captured the temporal alignment of global brain fluctuations, anti-correlation of the Default Mode (DMN)- and Task Positive (TPN)-like networks, and activity in neuromodulatory nuclei of the reticular formation. While visual stimulation could trigger a transient spatiotemporal pattern highly similar to intrinsic QPPs, global signal fluctuations and QPPs during rest periods could explain variance in the following visual responses. QPPs and the global signal thus appeared to capture a common arousal-related brain-state fluctuation, orchestrated through neuromodulation. Our findings provide new frontiers to understand the neural processes that shape functional brain states and modulate sensory input processing.

## Introduction

Resting state fMRI (rsfMRI) and task-evoked fMRI are powerful complementary techniques to study brain function (Bandettini, 2012; Fox and Raichle, 2007). The first investigates the intrinsically highly active nature of the brain, while the second studies the brain’s reflexive properties and less so considers the ‘background’ intrinsic fluctuations that are averaged out across trials (Raichle, 2010). Recent studies support the view that intrinsic BOLD fluctuations across individual trials affect sensory responses and behavioral performance (Boly et al., 2007; Fox et al., 2007, 2006; He, 2013; Sadaghiani et al., 2009; Thompson et al., 2013). Yet, it remains unclear which specific regional or brain-wide neural mechanisms underlie this interaction.

Answers may come from emerging tools in the field of time-resolved rsfMRI, which attempts to identify the dynamic interaction of brain networks during the resting state (Allen et al., 2014; Deco et al., 2011; Keilholz, 2014). Brain ‘states’ or cognitive fluctuations may be identified and their role in task performance evaluated (Gonzalez-Castillo et al., 2015; Keilholz et al., 2017; Kucyi et al., 2018). Changes in vigilance or attention may also be identified and appear difficult to dissociate from cognitive brain states (Allen et al., 2018; Chang et al., 2016; Hinz et al., 2019; Laumann et al., 2017; Shine et al., 2016; Tagliazucchi and Laufs, 2014; Wang et al., 2016).

One recurring finding is that whole-brain global BOLD signal dynamics contain an arousal component (Horovitz et al., 2008; Liu et al., 2017; Sämann et al., 2011; Wong et al., 2016, 2013, 2012; Yeo et al., 2015). A significant fraction of the global BOLD signal has been correlated with a global neuronal signal, which further appeared to be driven by cholinergic neuromodulatory actuators, and was coupled to arousal-related fluctuations in brain state (Chang et al., 2016; Liu et al., 2018; Schölvinck et al., 2010; Turchi et al., 2018; Wen and Liu, 2016). Despite these insights, there is currently a lack of well-defined brain state dynamics and associated properties.

New insights for further understanding the interplay of these processes across different brain areas and temporal lengths may come from recently developed techniques such as identifying and studying quasi-periodic patterns (QPPs) of brain activity. QPPs, first introduced by the Keilholz group in 2009 (Majeed et al., 2009), refer to infraslow (0.01-0.2Hz) spatiotemporal patterns in the BOLD signal that recur quasi-periodically throughout the duration of a resting state scan. Interestingly, across multiple species, QPPs display prominent anti-correlation between the Default Mode network (DMN) and Task Positive network (TPN) (A. Abbas et al., 2016; Belloy et al., 2018a; Majeed et al., 2011; Yousefi et al., 2018). The DMN and TPN are thought to regulate competing cognitive processes related to processing of internal and external input (Fransson, 2006; Greicius et al., 2003; Northoff et al., 2010). Fluctuations in their activity reflects modulations in attention, affects sensory responses, and can explain some behavioral variability (Abbas et al., 2019; Esterman et al., 2013; Helps et al., 2009; Lakatos et al., 2016; Sadaghiani et al., 2009; Weissman et al., 2006). Specifically, time-varying DMN-TPN anti-correlations have been correlated with arousal fluctuations and lapses in behavioral performance (A Abbas et al., 2016; Lynn et al., 2015; Thompson et al., 2013; Wang et al., 2016). Substantial evidence thus suggests that QPP dynamics reflect fluctuations in brain state and may modulate task-evoked sensory responses, yet this question has not been formally investigated.

These observations listed above suggest a functional overlap between global neural brain dynamics and QPPs, a link that has recently been supported through their spatiotemporal overlap (Belloy et al., 2018a; Nalci et al., 2017; Yousefi et al., 2018). This relationship is, however, not simple. On one hand, the ‘global’ signal may variably be composed by the activity of large resting state networks rather than brain-wide activations (Billings and Keilholz, 2018), while on the other hand, QPPs do not fully make up the global signal (Belloy et al., 2018b; Yousefi et al., 2018). The intricate relationship between the global signal and DMN-TPN anti-correlation has a longstanding history in the fMRI community (Murphy and Fox, 2016), but their mechanistic relationship remains to be identified.

In this study, we hypothesized that the quasi-periodic anti-correlations between the mouse DMN- and TPN-like networks, identified under the form of QPPs (Belloy et al., 2018a), may reflect ongoing brain state fluctuations linked to arousal- or salience-related processes. Further, we speculated that if this relationship between QPPs and brain state fluctuations exists, then this would establish an intricate and measurable link between QPPs and sensory response variance. To this end, we performed fMRI experiments in healthy C57BL6/J mice under rest and sensory visual stimulation conditions with two main goals: **1)** Determine how mouse resting state QPPs relate to global brain fluctuations, indicated to reflect arousal dynamics in the literature, and **2)** Determine if ongoing QPPs either affect, or are modulated by, visual sensory processing.

## Results

Experiments were performed in mice (N=24) that were further separated in two equally populated groups (N=12) that followed equivalent experimental procedures, resting state and visual fMRI, albeit with slightly different order to control for potential time and anesthesia effects **(Supplementary table S1**). Overall, scan mean frame-wise displacement was negligible across all scans [0.38 ± 0.04 mm (mean + STD)] (**Supplementary table S2)**. Resting state scans from multiple sessions and time points were used to determine large-scale resting state networks (RSNs), by means of ICA (**Supplementary Figure S1**). These RSNs displayed plausible physiological networks, supporting data quality (cfr. **Supplementary Text**). There were no significant differences of RSNs between animal groups (**Supplementary table S3**), nor significant differences of visual activation maps between groups, supporting data pooling for subsequent analyses.

### Quasi-periodicity during resting state

Using methods and analysis strategies that we previously established [cfr. (Belloy et al., 2018a, 2018b) and M&M], we consistently identified three QPPs of interest in the data (**Figure 1A**): QPP1, a short 3s pattern that displayed a transient wide-spread anti-correlation between DMN-like/Sensory networks and the lateral cortical network (LCN; a proposed mouse analogue of the TPN; cfr. discussion) (**Video 1**); QPP2, a 9s pattern that initially is similar to QPP1 but continues and reverses pattern in later frames (**Video 2**); and QPP3, a 9s pattern cycling between wide-spread activation and de-activation (**Video 3**). Except for the LCN, the three QPPs largely involved the same brain areas. All three QPPs displayed a high degree of temporal co-linearity (**Figure 1A-B**), indicating a potential shared underlying process. This was further exemplified by the common quasi-periodicity in their power spectra [see deviations from the 1/f power law; (**Supplementary Figure S2**)]. To further estimate the time-relationship of these three QPPs, phase-phase plots were constructed for all QPP pairs (**Figure 1C-E**). All QPPs displayed prominent phase-phase coupling, and this was only slightly reduced between QPP2 and QPP3. Notably, for each of the observed QPPs, an opposite phase variant was also observed with consistent temporal characteristics (**Supplementary Figure S3**). These were not further considered given their equivalence (nearly inverted time series) to the primary described QPPs.

**Figure 1.**
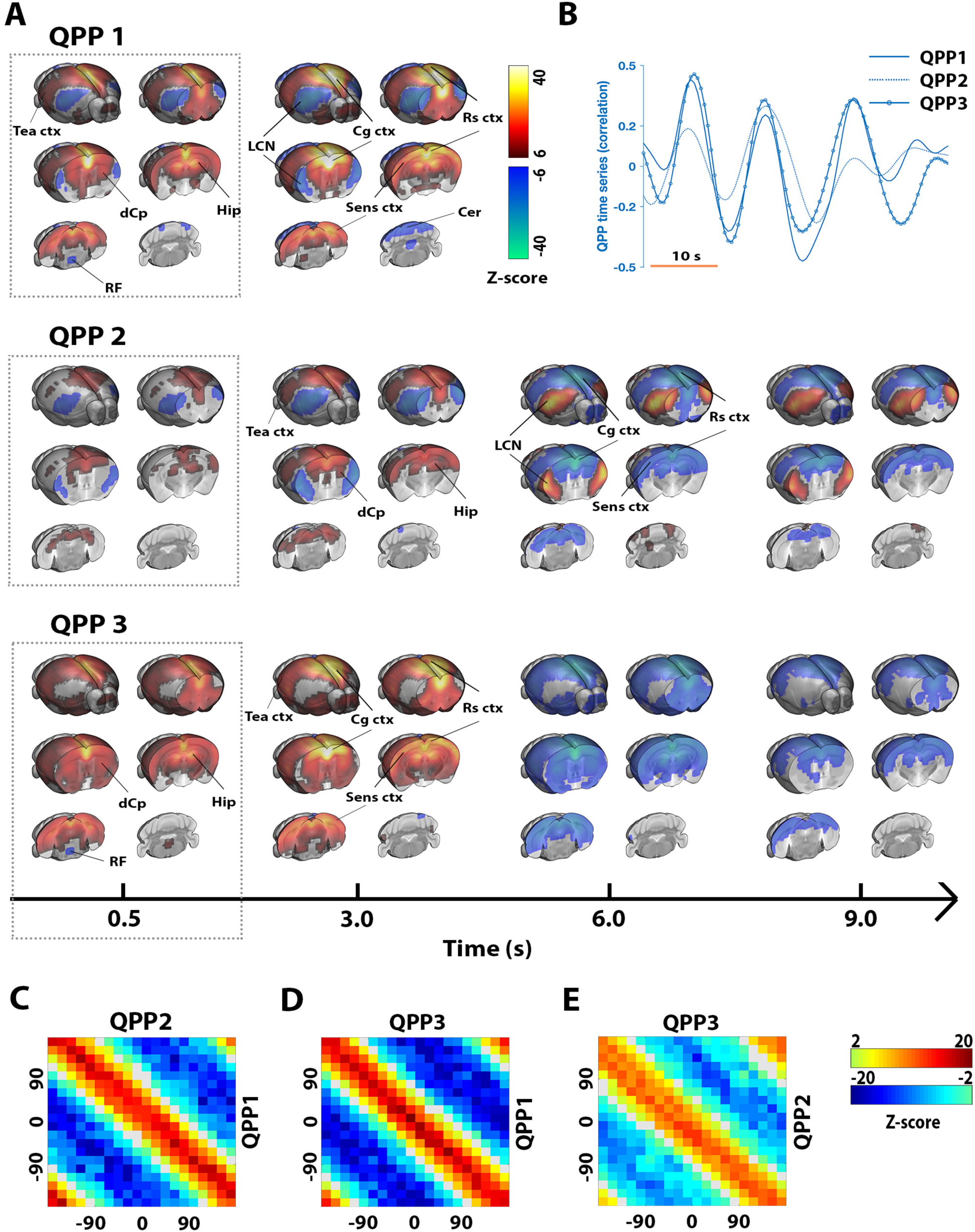
Three temporally co-linear quasi-periodic brain fluctuations identified during resting state. Three QPPs were identified **(A)**. QPP1 displayed a transient 3 s pattern of anti-correlation between DMN-like/Sensory networks and the LCN, QPP2 appeared similar as QPP1 but reverses in later frames, and QPP3 displayed cycling wide-spread activation and deactivation. Relevant brain areas are marked; DMN-like areas included Cg ctx, Rs ctx, Tea ctx, Hip, and dCp. The three QPPs displayed a high degree of co-linearity, evident both visually **(B)** and from phase-phase coupling **(C-E)**. **A-E)** n = 71 scans. **A)** QPPs are displayed on the same time axis [alignment through cross-correlation of QPP correlation vectors (B)]. Maps display Z-scores [Z-test with H0 through randomized image averaging (n=1000), FDR p<10^−7^, cluster-correction 4 voxels]. **B)** Single subject excerpt. QPP correlation vectors represent Pearson correlations of QPPs with functional image series. **C-E)** Phase-phase plots show Z-scores; center red diagonal marks strong co-phasic dynamics [first level Z-test with H0 through randomized circular shuffling (n=1000); second level Z-test, FDR p<0.05]. *Abbreviations. Quasi-periodic pattern, QPP; DMN, Default mode network; Lateral cortical network, LCN; Hippocampus, Hip; dorsal Caudate Putamen, dCp; Cingulate cortex, Cg ctx; Retrospleneal cortex, Rs ctx; Sensory cortex, Sens ctx; Cerebellum, Cer; Reticular formation, RF; False-discovery rate, FDR; repetition time, TR*.

### Co-linearity with global brain fluctuations

Given that DMN-TPN anti-correlations (reflected in the QPPs) and fluctuations in the global fMRI signal have independently been demonstrated to have relationships to sensory variance and/or changes in arousal, we next investigated the relationship between the global signal and QPPs. The global signal displayed wide-spread activations that strongly involved Sensory and DMN-like networks, followed by deactivation that was mainly confined to the Retrospleneal and Cingulate cortex. Interestingly, a focal brain stem deactivation at the level of the reticular formation was also observed during the widespread activations (**Figure 2A; Video 4**). Given recent findings indicating that neuromodulatory nuclei may regulate global signal fluctuations, this could suggest a potential mechanistic link (Liu et al., 2018; Turchi et al., 2018). Further, the global signal displayed marked temporal overlap with all three identified QPPs (**Figure 2B**) and a power spectrum similar to QPP1, but further reduced quasi-periodicity (Supplementary **Figure 2**). Phase-phase plots revealed that QPP3 was highly temporally co-linear with the global signal, followed by QPP1, which was also strongly co-linear with the global signal, and lastly QPP2, which displayed weaker phase-phase coupling (**Figure 2C-E**). In summary, the results presented in **Figure 1 & 2** suggested a slightly varying but consistent temporal alignment of all QPPs as well as the global signal, reinforcing their hypothesized link to a common underlying brain state process.

**Figure 2.**
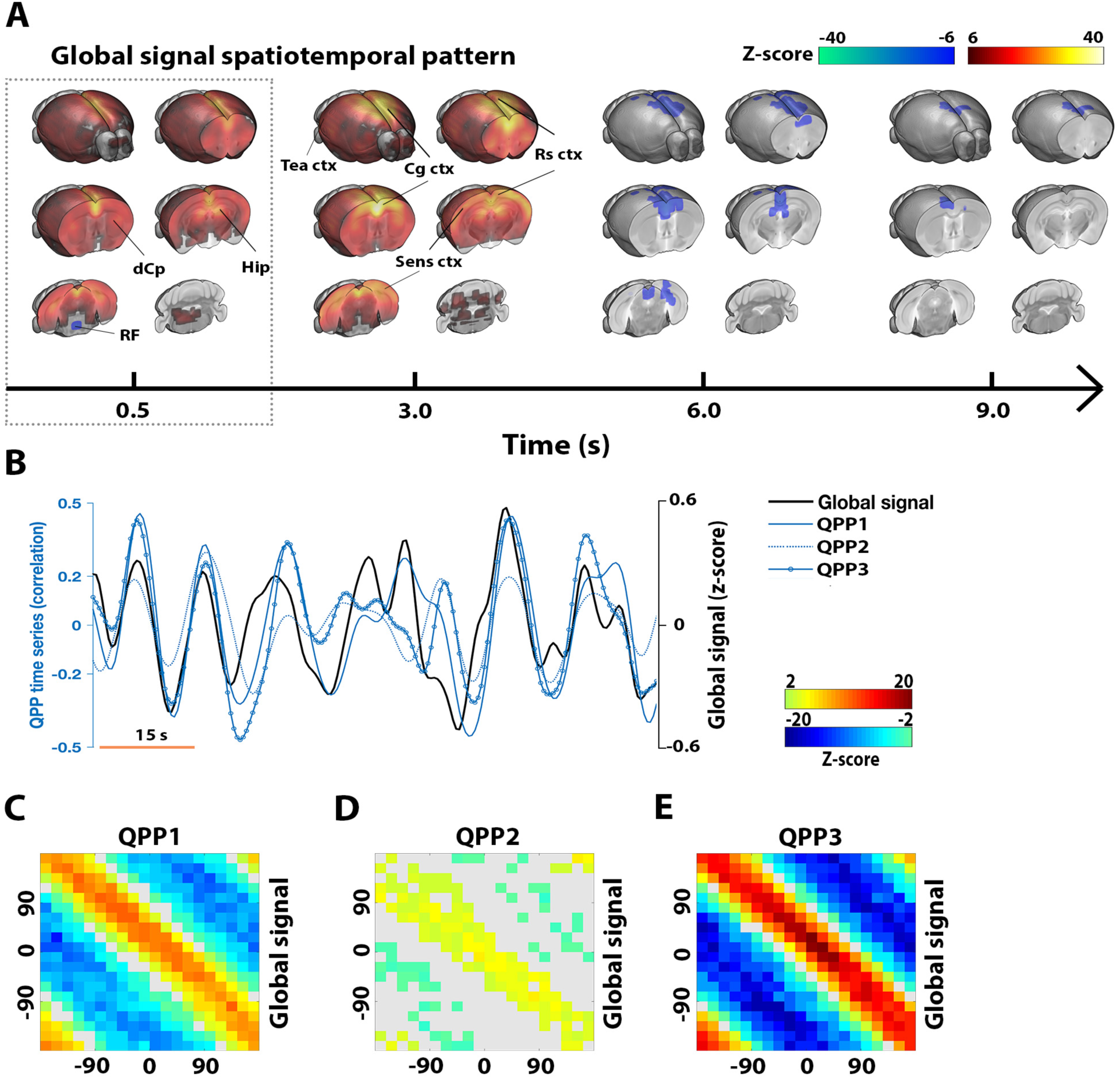
Quasi-periodic brain patterns temporally coincide with global brain fluctuations. The global signal was marked by a first phase of widespread activation, with stronger activations in sensory cortex and DMN-like areas **(A)**. A focal deactivation was also observed in the dorsal brain stem, at the height of the reticular formation. The second phase of the global signal mostly incorporated deactivation in Rs ctx and Cg ctx areas. Both visually **(B)** and based of phase-phase plots **(C-E)**, the global signal displayed clear temporal co-linearity with the three observed QPPs, with decreasing strength from QPP3 to QPP1 to QPP2. **A-E)** n = 71 scans. **A)** Maps display Z-scores [Z-test with H0 through randomized image averaging (n=1000), FDR p<10^−7^, cluster-correction 4 voxels]. **B)** Single subject excerpt. QPP correlation vectors represent Pearson correlations of QPPs with functional image series. Global signal fluctuations are shown as Z-scored BOLD intensities. Time series were aligned through cross-correlation (global signal peak occurred on average 2s into QPP1-3). **C-E)** Phase-phase plots show Z-scores [first level Z-test with H0 through randomized circular shuffling (n=1000); second level Z-test, FDR p<0.05]. *Abbreviations. Quasi-periodic pattern, QPP; DMN, Default mode network; Lateral cortical network, LCN; Hippocampus, Hip; dorsal Caudate Putamen, dCp; Cingulate cortex, Cg ctx; Retrospleneal cortex, Rs ctx; Sensory cortex, Sens ctx; Cerebellum, Cer; Reticular formation, RF; False-discovery rate, FDR; repetition time, TR;*

### Intrinsic brain response to visual stimulation

After determining the properties and temporal relationships of resting state QPPs and the global signal, we investigated if similar relationships can also be observed during a visual stimulus processing design that is expected to trigger changes in brain state. To this end, we used a visual stimulation block design (30s ON – 60s OFF) with intentionally long OFF periods to allow the activity to return to baseline each time before the next visual activation block. First, to identify the visually stimulated areas, we used a classical generalized linear model (GLM) approach by convolving the block-design paradigm with the hemodynamic response function (HRF) in order to derive the signal predictor (cfr. M&M). Clear activations were observed in areas related to visual processing: dorsal thalamic nuclei (including Lateral geniculate nucleus; LGN); Superior colliculus (S. Col), Visual cortex (Vis ctx) and Hippocampus (**Figure 3A**). Then, the QPP spatiotemporal pattern finding algorithm was used to determine if spatiotemporal patterns (STPs) similar to QPPs could be observed in the visual fMRI scans. In this case, in addition to the normal analysis, we also performed the STP estimation after performing global signal regression, which, we reasoned, could potentially remove brain wide responses induced by visual stimulation that would interfere with STP detection. Both with and without global signal regression, the resultant STPs were largely dominated by visual activations, and also brain-wide responses in prefrontal and lateral cortical areas, but they were not clearly reminiscent of resting state QPPs (**Supplementary Figure S4**).

**Figure 3.**
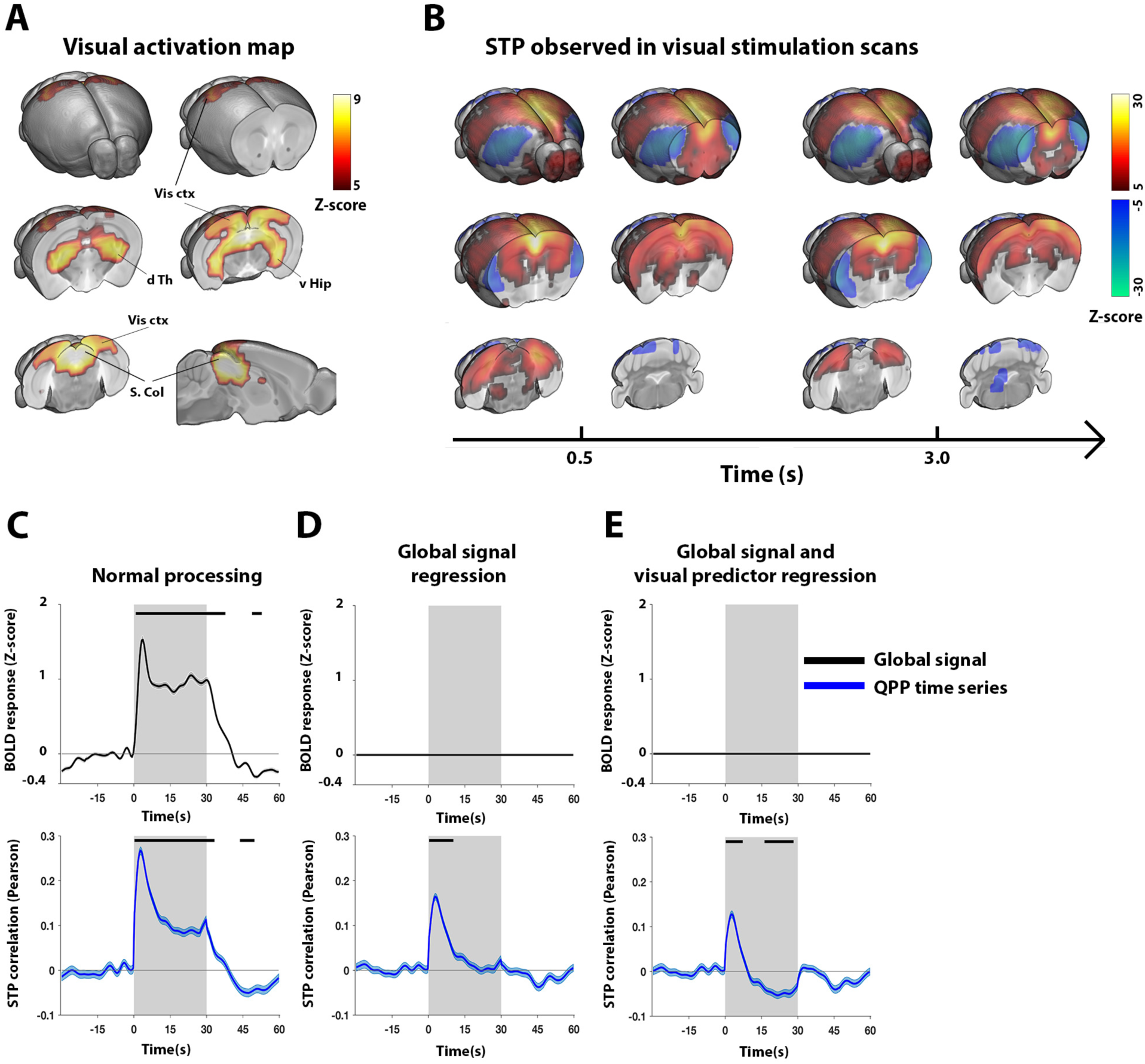
Visual stimulation intrinsically evokes a short quasi-periodic spatiotemporal pattern, also observed during resting state. Reliable visual activations were observed in brain areas related to visual sensory processing **(A)**. A short 3s STP, highly similar to QPP1 determined during resting state scans (spatial correlation = 0.90), was observed after regression of the visual predictor **(B)**. This task-derived STP displayed, on average, a peak correlation at the start of stimulation trials **(C-E)** and showed co-linear dynamics with the global signal **(C)**. The early peak correlation persisted even after regression of the visual and global signal **(E).** This short STP thus displayed co-linear, yet dissociable, response dynamics with visual activations, suggesting it represented an intrinsic response component rather than visual processing *per se*, nor was it solely the result of (a-specific) brain-wide activations. **A-E)** n = 24 scans. **A)** Maps display Z-scores (first level GLM; second level one sample T-test; T-scores normalized to Z-scores; FDR p<10^−5^, cluster-correction 4 voxels]. **B)** Maps display Z-scores [Z-test with H0 through randomized image averaging (n=1000), FDR p < 10^−5^, cluster-correction 4 voxels]. **C-E)** The global signal (top) and STP correlation vector (bottom), each respectively averaged across all trials and animals (n = 10 trials x 24 animals). Grey areas mark trials (ON periods), traces show mean (BOLD time courses demeaned and variance normalized to 10s OFF period prior to stimulation), patches show STE, black bars mark significance (one sample T-test, FDR p<10^−5^). *Abbreviations. Quasi-periodic pattern, QPP; Spatiotemporal pattern, STP; ventral Hippocampus, v Hip; dorsal Thalamus, d Th; Visual cortex, Vis ctx; Superior colliculus, S. Col; standard error, STE;*

To further eliminate STPs directly reflecting visual activation, we also performed the same analysis after the visual predictor was regressed from the task fMRI scans. Under these conditions, the spatiotemporal pattern finding algorithm revealed a short 3s STP that was highly similar to QPP1 during rest (spatial cross-correlation = 0.90; **Figure 3B**). Surprisingly, correlating this STP with the fMRI time series before regression of the visual or global signal predictors displayed, on average, a significantly increased correlation with the image series at the start of visual stimulation blocks (**Figure 3C**). No such response could be reliably observed for longer STPs (**Supplementary Text**; **Supplementary Figure S4 & S6**). Notably, the correlation increases of the 3s STP around the start of the visual stimulation were preserved after global signal regression (**Figure 3D**) as well as after regression of both the global signal and the visual stimulation predictors (**Figure 3E**). We therefore conjectured that this STP may represent an intrinsic component triggered by the visual stimulus but does not represent the visual sensory processing per se. This result is further supported by the higher spatial correlation of this STP with the resting state QPP in comparison to the visual activation profile (spatial cross-correlation = 0.56 when excluding significantly activated areas [cfr. **Figure 3A**]) and, in addition, by the fact that this STP was also observed at different time-points beyond the start of the visual stimulation blocks, such as during off periods and occasionally at different phases of the visual stimulus (**Supplementary Figure S5**). These results thus suggest that the observed STP represents a default ongoing brain fluctuation that can be modulated by visual stimulation.

### Intrinsic quasi-periodicity explains visual response variance

In the previous section, we demonstrated that visual stimulation can modulate, beyond sensory processing, intrinsic brain dynamics as reflected in STPs. Here, we asked the question if intrinsic brain dynamics could also influence sensory responses. To this end, we investigated whether signal fluctuations in visual areas prior to visual stimulation (stimulus OFF interval), could explain a portion of the visual response variance during stimulation (**Figure 4A**). Across animals and trials, signals in time bins prior to stimulation were correlated with those in the time bin of the visual response peak. The prior time point with the highest correlation value (essentially the one with the highest predictive power) was used to stratify stimulation trials into: high pre-stimulus (HP), medial pre-stimulus (MP), and low pre-stimulus (LP) visual signal amplitudes. HP trials displayed significantly higher peak and plateau responses compared to MP trials, while LP trials displayed significantly lower peak responses compared to HP trials (**Figure 4B**). To gain a mechanistic understanding of these observations, the same trial sets were evaluated under conditions of global signal and STP regression (**Figure 4C-E**). Differences in visual responses were still apparent under conditions of STP regression but became less pronounced (**Figure 4B-D**). After global signal regression, or combined STP and global signal regression (**Figure 4C&E**), no more significant differences could be observed between stratified trial sets. The absolute amplitudes of HP and LP visual signals (prior peaks or dips) decreased respectively by 64% and 49% after STP regression. With global signal regression, inversions were observed for HP and LP signals. After global signal regression, absolute amplitudes decreased by respectively 78% and 56%, and, after combined STP and global signal regression, by 56% and 71%. Notably, across the stimulus duration, only combined global signal and STP regression reduced the variance of the visual signal to levels almost equal to those observed during stable rest periods (**Supplementary Figure S7**).

**Figure 4.**
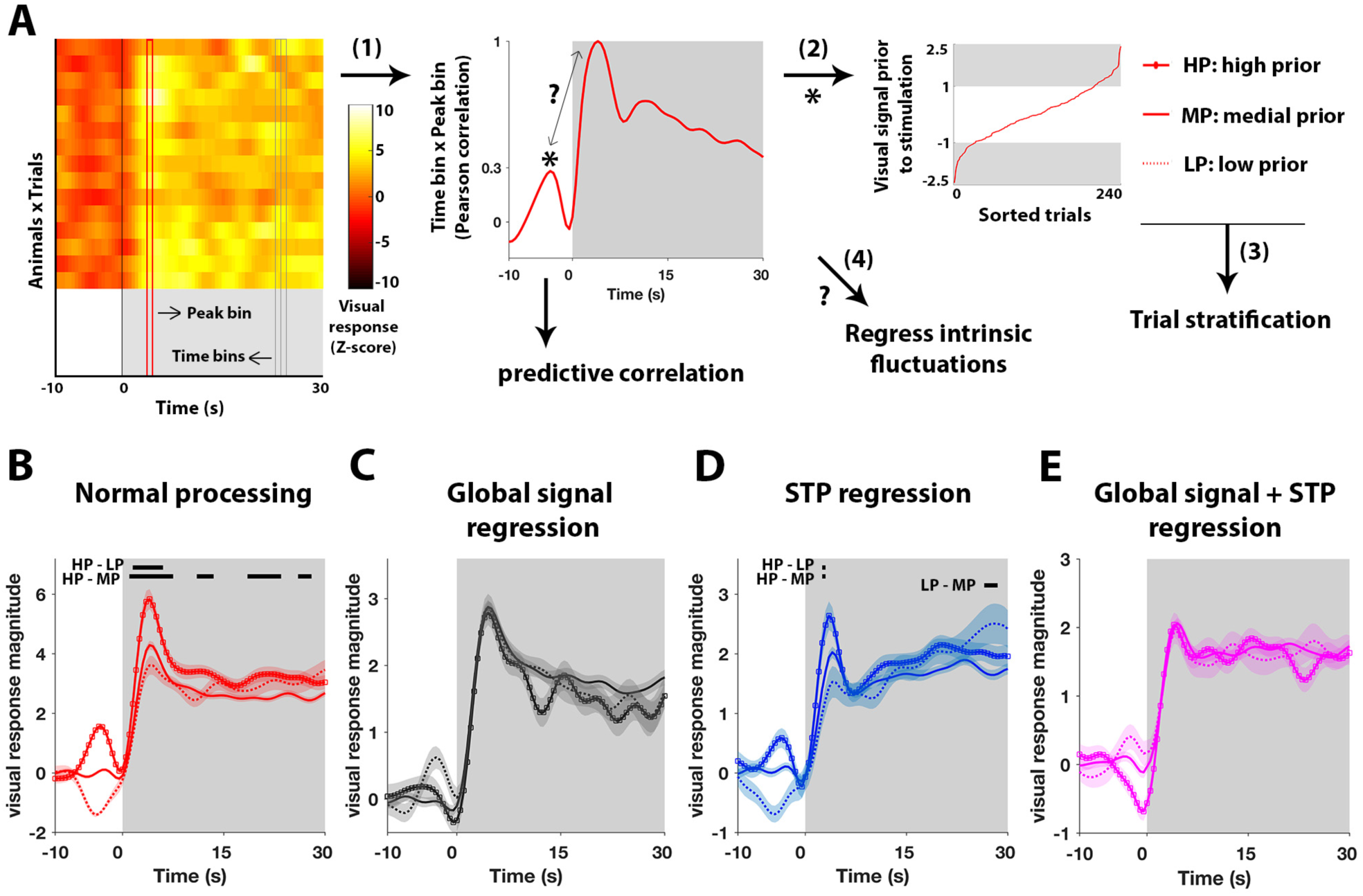
Intrinsic brain-wide quasi-periodicity predicts visual response variance. Stimulation trials were stratified into sets based on intensities in visual areas prior to stimulation: high prior (HP), medial prior (MP), and low prior (LP) trials **(A)**. With normal processing, clear differences were apparent between trial sets, particularly for the initial peak response (**B**). For the same trial sets, after STP regression, differences were diminished **(D)**, while after either global signal **(C)** or STP + global signal regression **(E)**, no more differences were observed. **A-E)** n = 24 animals x 10 trials. Time traces are demeaned and variance normalized to 10s OFF period prior to stimulation. **A)** Illustration of individual stimulation trials, time bins (grey) and response peak bin (red). (1) Identifying maximal correlation (*) of time bins prior to stimulation with response peak. (2) Sorting of visual signal intensities prior to stimulation (grey patches > 1 STE). (3) Stratification. (4) Evaluating role of intrinsic brain dynamics through regression analyses. **B-E)** Black bars indicate significant mean differences between trial sets (One-way ANOVA, FDR (#bins) p<0.05; post hoc Bonferroni correction). *Abbreviations. Spatiotemporal pattern, STP; False-discovery rate, FDR; standard error, STE; analysis of variance, ANOVA*.

### Co-linearity with fluctuations in the reticular formation

Our results of the resting state data indicated collinearity of intrinsic brain fluctuations (QPPs and global signal) suggesting a potential link to an underlying process related to brain state. Similarly, analysis of visual stimulation scans demonstrated interactions between sensory processing and intrinsic brain fluctuations, indicating that these processes are finely intertwined. Interestingly, detailed observation of the QPPs and the global signal pattern unveiled that a focal area at the dorsal part of the brain stem cycled antagonistically with overall brain-wide activity (**Figure 5A**). To identify the cytoarchitectonic location of this area, we co-registered the MRI data to the Allen mouse brain atlas. This revealed that this area contained mainly pontine nuclei of the reticular formation (RF; **Figure 5B**). The average RF time courses across all three QPPs and the global signal were highly similar (**Figure 5C**), with an initial significant dip, followed by a significant peak approximately 4.5s later. Furthermore, to understand if this area could be related to intrinsic brain fluctuations during visual stimulation, we plotted the average initial time frames of the event-related activation maps (**Figure 5D**). Surprisingly, significant de-activations in the RF were observed time-locked to the start of visual stimulation. The time course of RF activity is presented in **Figure 5E**.

**Figure 5.**
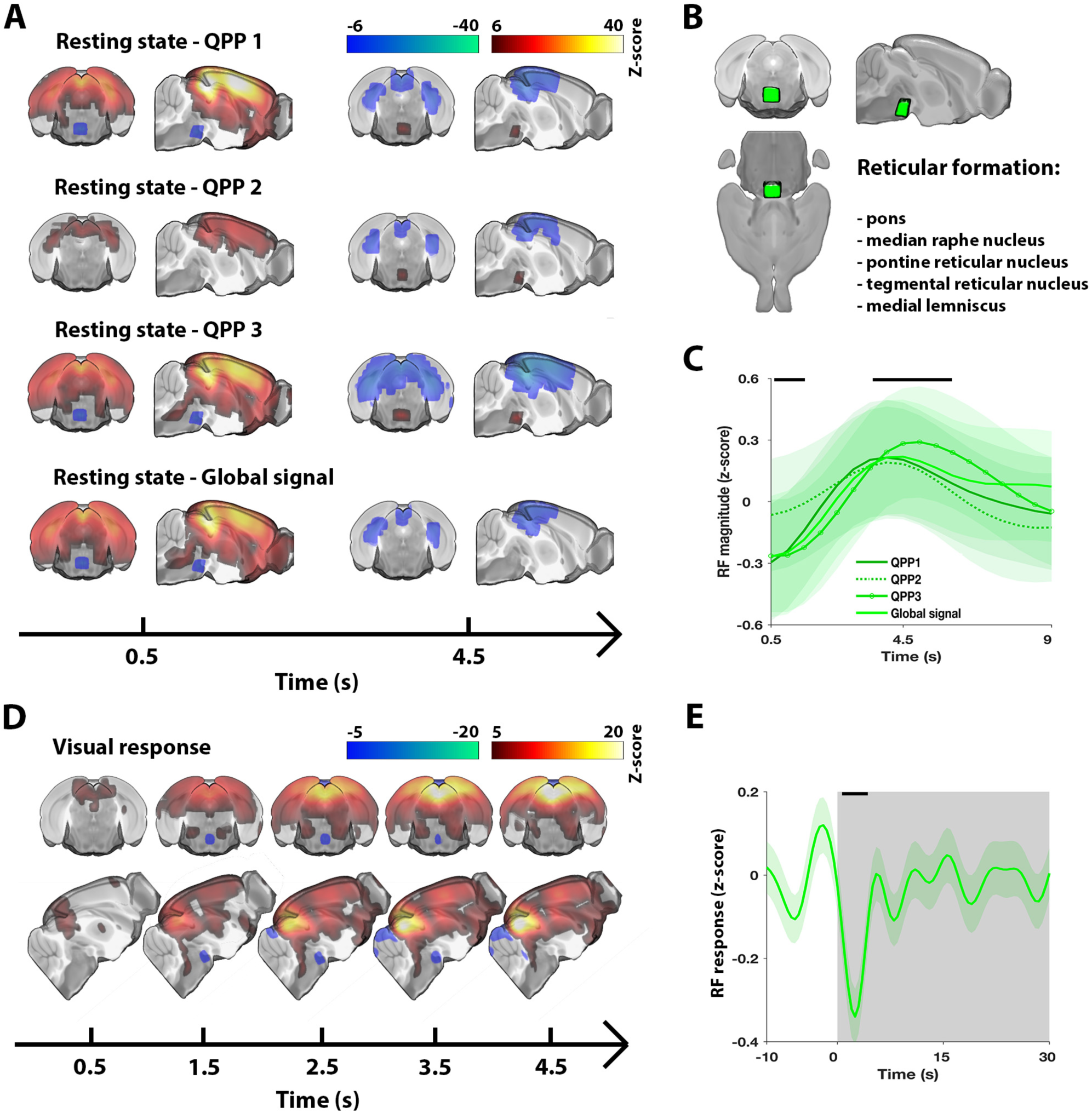
Activity in the reticular formation couples with quasi-periodic brain dynamics across the rest/task spectrum. All QPPs and the global signal displayed significant activity in a focal dorsal brain stem area. Anatomical labelling through co-registration with the Allen mouse brain atlas highlighted that this area contained nuclei of the reticular formation **(B)**. Time courses of the RF were on average highly similar across investigated spatiotemporal patterns **(C)**. The RF also displayed de-activation at the start of stimulation blocks **(D-E)**. **A)** n = 71 scans. Maps display Z-scores [Z-test with H0 through randomized image averaging (n=1000), FDR p<10^−7^, cluster-correction 4 voxels]. **B)** Visual rendering of focal brain area observed in (A) and (D). List indicates anatomical structures contained within this area. **C)** n = 71 scans. Average RF time series across respective QPP correlation or global signal peaks (traces show mean; patches show STE). Black bars mark significant deviation from zero [statistical test as in (A)]. **D)** Visual response averaged across trials and animals (n = 10 trials x 24 animals) for first 5s of stimulation. Voxel-wise time courses were demeaned and variance normalized to 10s OFF period prior to stimulation. Maps display Z-scores [one sample T-test; T-scores normalized to Z-scores; FDR p<10^−5^, cluster-correction 4 voxels]. **E)** Grey areas mark trials (ON periods), trace shows visual area signal mean, patch shows STE, black bar marks significance (one sample T-test, FDR p<10^−5^). *Abbreviations. Quasi-periodic pattern, QPP; False-discovery rate, FDR; standard error, STE*.

## Discussion

Many questions remain on the mechanisms through which intrinsic brain dynamics and sensory processing interact. We sought answers using fMRI in lightly anesthetized mice to track spatiotemporal activity patterns at the whole brain level, an approach that may provide new insights in comparison to more commonly performed invasive single site recordings. A vast emerging literature suggests that intrinsic global signal fluctuations, DMN-TPN anticorrelations, arousal dynamics, and neuromodulation, may all share common ground and could affect sensory processing. Our results provide evidence that quasi-periodic patterns captured an overall temporal alignment between these related phenomena. We further showed that, with high probability, visual stimulation evoked a spatiotemporal pattern highly similar to QPPs, with persevered co-linearity to the brain global signal and deactivations in the reticular formation. Finally, we showed that QPPs and the global signal could significantly predict a portion of the visual response variance. In summary, our findings suggest that QPPs and the global signal in mice likely capture a single brain-state fluctuation, mechanically coupled through neuromodulation, and we provide evidence that these spatiotemporal patterns affect sensory response variance.

QPPs observed here were highly consistent with those observed in previous mouse studies using single slice recordings (Belloy et al., 2018b, 2018a). Specifically, QPPs displayed widespread anti-correlation between the commonly observed mouse LCN and DMN-like/sensory networks (Grandjean et al., 2017; Liska et al., 2015; Zerbi et al., 2015). No direct evidence has so far been presented to identify a mouse TPN-like network, but the LCN has been suggested as the most likely candidate (Liska et al., 2015; Zerbi et al., 2015). This is further supported by the LCN’s anti-correlation with the DMN-like network, both in conventional functional connectivity analysis and within QPPs, highlighting consistency with human resting state network properties. We therefore discuss the LCN interchangeably with “mouse TPN-like network’. Further, the three identified QPPs displayed a high degree of temporal co-linearity, suggesting they likely reflected variants in a single spatiotemporal pattern. One possibility is that the shorter QPP1 was more likely to occur (stronger correlation vector) while the longer QPP2 (weaker correlation vector) identified instances where QPP1 oscillated and reversed in later frames. Alternatively, it is possible that due to temporal collinearity with brain-wide (de-)activations (i.e. the global signal and QPP3), the spatiotemporal pattern finding algorithm would have been biased towards lower correlation amplitudes for QPP2. The infraslow network dynamics observed here within QPPs are consistent with the quasi-oscillatory dynamics of co-activation patterns (CAPs, i.e. instantaneous brain activity patterns) previously observed in humans and mice (Gutierrez-Barragan et al., 2018; Liu and Duyn, 2013). While CAPs identified a richer set of dynamic network topologies, the QPP approach identified the most dominantly recurring brain-wide spatiotemporal pattern that likely comprised several temporally aligned CAPs. Notably, QPP1 displayed diminished periodicity compared to QPP2/3. This could suggest that shorter QPPs observed here more closely resemble 1/f aperiodic brain dynamics (He and Raichle, 2009). However, short QPPs have been shown to extend to variable-length QPPs (Belloy et al., 2018a). The signal of QPP1 is thus comprised of several band-limited oscillations (i.e. it displays a scale-free autocorrelation profile), which can give rise to an arrhythmic power spectrum (Palva and Palva, 2012).

The global signal spatiotemporal pattern displayed wide-spread activations, with stronger focal increases in sensory cortex and core DMN-like areas such as the dCP, dTh, dHip, Cg and Rs cortex. Limited (quasi-)periodicity was observed in the global signal’s temporal structure. This is in line with prior human studies that indicated the DMN-like network and sensory cortex as strong contributors to the global signal, which displayed only faint periodicity (Billings and Keilholz, 2018; Fox et al., 2009). We observed here that the global signal and QPPs displayed strong temporal collinearity. At the same time, QPPs/STPs that displayed regional anti-correlation could still be detected after global signal regression in both resting state and visual fMRI scans. The latter is consistent with prior work on QPPs (Belloy et al., 2018a; Majeed et al., 2011; Yousefi et al., 2018), and is reminiscent of the relationship between DMN-TPN anticorrelation and global signal regression (Fox et al., 2009; Murphy and Fox, 2016). Specifically, estimation and regression of QPPs and the global signal are qualitatively distinct (Billings and Keilholz, 2018). On one hand, global signal regression zero centers the instantaneous distribution of brain intensities, thereby preserving time-varying inter-regional variation. Even when strongly co-linear, global signal regression cannot fully remove QPPs with regional anti-correlation. On the other hand, QPP time courses reflect time-varying image similarities to a recurrent spatiotemporal template, which contains both global and regional variation. Inherently, some overlap between QPPs and the global signal is thus expected, but the extent of temporal alignment that was observed in this study, and the specific involvement of major resting state networks, are striking and suggestive of a shared physiological substrate. Another resting state study in mice, using different anesthesia, also observed strong phase coupling between the global signal and oscillatory activation patterns similar to QPPs described here (Gutierrez-Barragan et al., 2018), suggesting our findings can be generalized across mouse studies.

Activation maps in response to visual stimulation were highly consistent with those previously reported in mice (Niranjan et al., 2016). Visual responses displayed fast peak activations followed by stable plateau periods, consistent with fast haemodynamics in the mouse brain (Drew et al., 2011; Pisauro et al., 2013). Most mouse fMRI studies to date have focussed primarily on somatosensory stimulation paradigms, reporting strong variability in evoked responses and a-specific brain-wide activations on top of somatosensory networks responses (Adamczak et al., 2010; Reimann et al., 2018; Schlegel et al., 2015; Schroeter et al., 2016, 2014). These studies indicated that part of the brain-wide responses was due to transient increases in mean arterial blood pressure, caused by the arousal-promoting noxious nature of presented stimuli. In pilot studies, we did not clearly observe such responses for our visual stimulation and anesthesia protocols. Both QPPs and the global signal have furthermore been related to a neuronal substrate (Grooms et al., 2017; Pan et al., 2013; Schölvinck et al., 2010), while no clear link between QPPs and physiology could be established in prior mouse work (Belloy et al., 2018a). We thus propose that global signal and STP dynamics in response to visual stimulation may indeed reflect an arousal-related response, but one that is more likely of neuronal origin (see further below).

The visual-evoked STP indicated deactivation of the TPN-like network and activation of the DMN-like/sensory networks. This apparent task-related DMN activation may be considered counter-intuitive with regard to conventional observations that task engagement causes decreased DMN activity and increased TPN activity (Fransson, 2006; Northoff et al., 2010). Similarly, we observed that DMN activity in QPPs and the global signal correlated with larger visual responses, while some studies related DMN activity to decreases in sensory responses and increased response times (Helps et al., 2009; Weissman et al., 2006). In contrast, other studies reported a less canonical role of the DMN that is more consistent with the current findings (Esterman et al., 2013; Kucyi et al., 2017, 2016; Sadaghiani et al., 2009). In the latter, DMN activity reflected an attentive state, while TPN activity was associated with increased behavioral variance and suppressed attention. For instance, DMN and TPN activity just prior to auditory stimuli correlated respectively with significant increases and decreases in stimulus perception hit rate (Sadaghiani et al., 2009). This is consistent with our finding that a visual signal peak four-to-three seconds prior to stimulation could predict larger visual responses, but that both the prior visual signal amplitude and response variance were reduced after QPP and global signal regression. Some of the intrinsic self-predictive power of brain areas observed here, and in prior studies, may therefore be attributable to the ongoing anti-correlations between the DMN and TPN. This hypothesis was formally proposed in prior work, suggesting that global rhythmic anti-correlations of the DMN and TPN cycle the brain state between attentional lapses and periods of improved sensory entrainment (Lakatos et al., 2016). Currently, it remains unclear into what extent DMN- and TPN-like task dynamics in mice would be comparable to those in humans. Under anesthetized conditions, it is less likely that DMN/TPN dynamics would actually reflect human canonical responses to a cognitive challenge. It thus seems more likely that QPPs and the evoked STP in fact reflect a brain state dynamic with distinct physiological and arousal-related properties.

In addition to the visually-evoked STP, we also observed a temporally co-linear global brain response during stimulation. The global signal displayed strong predictive power for visual responses, while global signal regression reduced visual response variance. In agreement, several studies have shown that global brain fluctuations, and the reflected changes in global brain state, can modulate sensory responses (Lee and Dan, 2012; Mcginley et al., 2015; Pisauro et al., 2016; Schölvinck et al., 2015; Schroeder and Lakatos, 2010). In mice, global haemodynamic fluctuations were corelated to fluctuations in arousal state and superimposed on local neuronal processing of visual input (Pisauro et al., 2016). In cats, during rest periods and in response to visual stimulation, global fluctuations underlied a high degree of shared variance across primary visual cortex neurons (Schölvinck et al., 2015). After global signal regression, the inter-trial variability in visual responses could be reduced in a similar fashion to what we observed here for mouse BOLD responses. Our findings thus strengthen the emerging concept that, in addition to noise components, global signal fluctuations also reflect arousal fluctuations (Liu et al., 2017).

A consistent observation across all spatiotemporal patterns was the co-linear activity in a focal brain stem area that comprised brainstem nuclei of the reticular formation. This may provide some mechanistic understanding for the arousal-related phenomena seen in this study. The ascending reticular activating system (comprising the RF) is responsible for promoting wakefulness and attention through the orchestrated activity of neuromodulatory nuclei, such as raphe nucleus, locus coeruleus and nucleus basalis. Liu et al. (2018) showed that the global signal coincides with deactivation in the nucleus basalis in humans, while Turchi et al. (2018) could supress global signal fluctuations by directly inactivating this cholinergic nucleus in macaques. Addtionally, optogenetic activation of the serotonergic dorsal raphe nucleus in mice caused widespread deactivation of DMN-like areas (Grandjean et al., 2019), which reflected the spatiotemporal patterns observed in our study. Further, neuromodulatory structures are natural rhythm generators that provide infraslow patterned input to the brain (Drew et al., 2008). Different nuclei in humans have been functionally connected to the DMN (dorsal raphe nucleus) and TPN (locus coeruleus) (Bär et al., 2016). This could help reconcile the co-linear dynamics between QPPs and the global signal, which may arise due to the complex interplay of subcortical nuclei. Finally, neuromodulation can adaptively affect brain states to modulate processing of sensory stimuli (Lee and Dan, 2012; Safaai et al., 2015), which could explain transient deactivation of the RF in response to visual stimulation (additional discussion in **Supplementary Text**). Future experiments will be required to tease out the potential neuromodulatory regulation of QPPs, the global signal, and neuronal circuit structure of arousal in the mouse brain, using tools such as optogenetics and pupil-tracking (Carter et al., 2010; Joshi et al., 2016; Reimer et al., 2014).

In summary, this study provides insights into the mechanisms that couple resting state dynamics to sensory processing and points out research avenues to elucidate their underlying neural substrate. Our work is directly relevant for other pre-clinical studies in rodent models that likely face some of the intrinsic sensory response variability highlighted here. Lastly, our analytical approach may help increase understanding of neurological disorders in which neuromodulation and arousal are pertinent.

## Material and Methods

### Ethical statement

All procedures were performed in strict accordance with the European Directive 2010/63/EU on the protection of animals used for scientific purposes. The protocols were approved by the Committee on Animal Care and Use at the University of Antwerp, Belgium (permit number 2017-38), and all efforts were made to minimize animal suffering.

### Animals

MRI procedures were performed on 24 male C57BL/6J mice (Charles River) between 18 and 22 weeks old. Animals were first anesthetized with 3.5% isoflurane and prepared in the scanner according to routine practice (details in **Supplementary methods**). For functional scans, animals were anesthetized with a 0.075mg/kg bolus subcutaneous injection of medetomidine (Domitor, Pfizer, Karlsruhe, Germany), after which isoflurane was gradually lowered to 0.5% over the course of 20min. A subcutaneous catheter allowed continuous infusion of 0.15mg/kg/h medetomidine starting 15min post-bolus. This anesthesia regime is similar to an established optimal light anesthesia protocol for mouse rsfMRI (Belloy et al., 2018a; Grandjean et al., 2014). Acquisition of functional scans started 30min post-bolus. Physiological parameters were monitored for stability throughout scan sessions. Animals were scanned twice, two weeks apart (**Supplementary Table 1**).

### MRI procedures and registration

MRI scans were acquired on a 9.4T Biospec system (Bruker), with a four-element receive-only phase array coil and volume resonator for transmission. Briefly, anatomical scans were acquired in three orthogonal directions to render slice position consistent across animals. Initial fMRI scans lasted 10min, and directly following fMRI scans (rest or visual stimulation) lasted 15min. In each session a 3D anatomical scan was also acquired. The open source registration toolkit Advanced Normalization Tools (ANTs) was used to construct a study-based 3D anatomical template. The study EPI template was then registered, in a 2-stage procedure, to the Allen brain mouse atlas (Oh et al., 2014). Further presented analysis of functional EPI data was thereby kept within the EPI template space. Additional details are provided in **Supplementary Methods**.

### Visual stimulation design

Bin-ocular visual stimulation with flickering light (4Hz, 20% duty cycle) was presented to the animals by means of a fiber-optic coupled to a white LED, controlled by a digital voltage-gated device (Max-Planck Institute for Biological Cybernetics, Tuebingen, Germany) and a RZ2 bioamp processor (Tucker-davis technologies). Stimulation paradigms were triggered by a TTL pulse output from the scanner at the beginning of the EPI sequence. Visual stimulation scans lasted 15 min and visual stimuli were presented in a block design: 30s ON, 60s OFF, repeated 10 times with the first stimulus starting 30s post scan start.

### Functional scan pre-processing

Motion parameters were obtained for each scan (six rigid body transformation parameters), images were realigned and normalized to the study-based mean EPI template and smoothed (σ = 2 pixels) [Statistical Parametric Mapping (SPM12) software (Wellcome Department of Cognitive Neurology, London, UK); MATLAB2017b]. Motion parameters were regressed out of the fMRI scans and images were filtered using a 0.008-0.2Hz butterworth IIR filter, detrended, demeaned and normalized to unit variance (z-score operation). For visual-evoked fMRI scans, demeaning and variance normalization was performed with regard to 10s OFF periods prior to stimulation (z-scoring procedure: Z = (*x* − *μ*)/σ, with x = sample, μ = sample mean, σ = sample standard deviation). Time points at start and end of the image series were removed to account for filtering effects. Depending on the desired analysis, global signal regression (GSR) was performed. To determine spatiotemporal patterns, a brain mask was used to exclude ventricles.

### Spatiotemporal pattern finding algorithm

QPPs/STPs were determined using the spatiotemporal pattern finding algorithm described by Majeed and colleagues in 2011 (Majeed et al., 2011). Shortly, the algorithm identifies BOLD spatiotemporal patterns (distribution and propagation of BOLD activity across different brain areas over the duration of a specific predefined time-window) that recur frequently over the duration of the functional scans. The process is unsupervised and starts by randomly selecting a starting template from consecutive frames in the image series, corresponding to the predefined time-window length. Then, this template is compared with the image series via sliding template correlation (STC). A heuristic correlation threshold (ρ>0.1 for the first three iterations and ρ>0.2 for the rest) is used to define sets of images at peak threshold crossings that are averaged into a new template. This process is repeated until convergence. As the outcome of this procedure depends on the initial, randomly selected starting pattern, the process was repeated multiple times (n = 250) with randomly selected seed patterns from different time-points in the time-series. The process was also repeated for multiple window lengths (3-12s, 1.5s intersperse) as STP length is not known *a* priori. QPPs were obtained by applying the algorithm to the concatenated time series of all individual subjects within a group. Detailed descriptions of the algorithm, and videographic illustrations, are provided elsewhere (Belloy et al., 2018a; Majeed et al., 2011).

### Quasi-periodic pattern selection

After the spatiotemporal pattern finding algorithm concluded identifying the large set (n = 250 x 7 window sizes) of possible patterns, we proceeded to identify the patterns of interest based on prior knowledge, their similarity, and their STCs (herein often referred to as QPP time series) that indicate occurrences (correlation peaks) and time-varying similarity to the functional scans. It was previously established that both short (3s) and long (9s) QPPs can be uniquely identified from mouse (Belloy et al., 2018a, 2018b), and rat (Majeed et al., 2011), rsfMRI recordings. In these studies, short 3s QPPs displayed the strongest time-varying correlation and were always marked by spatial anti-correlation of various brain areas, while longer QPPs displayed lower amplitude time-varying correlation, could also display brain-wide activity, and tended to capture bi-phasic extensions of shorter QPPs. Given these known priors, we opted to first identify 3s QPPs. Then, QPPs were also defined for other window sizes. Specifically, for each window size, we selected as the most representative QPP the one that displayed the highest sum of correlation values at QPP occurrences [cfr. (Yousefi et al., 2018)]. From the resultant set of QPPs, the window size corresponding to a full cycle bi-phasic pattern was calculated [cfr.(Belloy et al., 2018a)]. All analyses were performed with and without global signal regression; findings for both approaches were integrated (cfr. below). Additional details are provided in **Supplementary Methods**.

### Significance maps

The number of QPP occurrences (ρ>0.2 threshold crossings) decreases with longer window sizes. Further, QPPs were determined with and without global signal regression. Therefore, to aid QPP comparisons, a homogenization procedure was employed. QPPs determined after global signal regression, were correlated with image series for which no global signal regression was performed. The resultant correlation vector was used to calculate QPP occurrences. Further, after QPPs were defined, the correlation threshold (ρ>0.2) was reduced for longer QPPs so that an equal number of occurrences was achieved as for short 3s QPPs. For each QPP, significant voxels were defined from each voxel’s intensity distribution of unique image frames contained within the QPP. This was evaluated for each QPP time frame respectively and through H0 estimation. Specifically, for each respective voxel and time frame within a QPP, a T-score was calculated (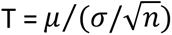) for its distribution of signal intensities (μ = mean; σ = standard deviation, n = sample size). For an equal n, 1000 reference distributions were calculated through randomized image frame selection. For each reference, a respective T-value was determined to construct the H0 distribution. A Z-test was employed to evaluate significance. Resultant significance maps were false discovery rate (FDR)- and cluster-size corrected (threshold = 4 voxels).

To visualize the global signal, image frames surrounding global signal peaks were averaged into a spatiotemporal template, i.e. a global signal co-activation pattern (CAP). This approach is consistent with the methodology presented by Liu and Duyn (Liu and Duyn, 2013), but includes temporal extension of signal peaks. A detailed description of this method is described elsewhere (Belloy et al., 2018a). An activation map of the global CAP, and related statistical analysis, was calculated in the same way as described for QPPs (cfr. above).

As a final homogenization step additional image frames that followed the core of short QPPs (e.g. 3s) were included to allow comparison with other longer spatiotemporal patterns. In this procedure, there is no re-estimation of the QPP or its correlation vector, only additional image frames following correlation peaks are averaged into the elongated template.

For each visual fMRI scan, the stimulation paradigm was convolved with a haemodynamic response function (HRF). The resultant visual predictor was used within a generalized linear model (GLM), i.e. first-level analysis, to derive subject voxel-wise parameter coefficients (β) and T-values. Subject activation T-maps were then evaluated at the group level, i.e. second-level analysis, by means of a one-sample T-test (H0 distribution: μ = 0 and σ = sample σ). Resultant group average activation maps were FDR- and cluster-size corrected (threshold = 4 voxels). The HRF was based on a literature-driven ground truth estimate (details in **Supplementary Methods**). Further, time-frame by time-frame group-average visual activation maps were also constructed by analyzing voxel-wise intensity distributions at each time point across all trials and animals (n = 24 animals x 10 trials). One-sample T-tests were used to define significant (de-)activations (H0 distribution: μ = 0 and σ = sample σ). Resultant frame-wise group-average activation maps were FDR- and cluster-size corrected (threshold = 4 voxels). For consistency, T-scores were standardized to Z-scores using the normal cumulative distribution function.

### Phase-phase coupling

Contrary to conventional correlation-based approaches, phase-phase coupling analysis can provide more detailed information regarding the relationship of two signals. Particularly, it can be used to calculate whether signals display in-phase, out-of-phase, or anti-phase properties, while no assumptions are made about causality or directionality. Prior work established that phase estimation for QPPs from rat rsfMRI data is feasible (Thompson et al., 2014), as well as for global signal and network fluctuations from mouse rsfMRI (Gutierrez-Barragan et al., 2018). Thus, for each subject respectively, the instantaneous phase of QPP or global signal time series were extracted using the Hilbert transform. Phase data was then binned across the [−π, π] range and the number of matching observations between two respective signals were counted on phase-phase grids (normalized to scan length). For each subject, an H0 distribution was obtained by randomly (n=1000) shifting one of two time-courses forward or backward in time [−10s:0.5s:10s] and filling in the phase-phase grid at each instance. For each voxel in the grid, the real value was evaluated with regard to the normal H0 distribution and a Z-score was derived. This procedure was repeated for all subjects, so that each voxel within the group-level phase-phase grid contained a distribution of Z-scores. For each voxel on the group grid, one-sample T-tests were used to define significant deviations from zero (H0 distribution: μ = 0 and σ = sample σ). The resultant significance map was FDR-corrected. For consistency, T-scores were standardized to Z-scores using the normal cumulative distribution function.

### Regression and visual response analyses

Various multiple linear regression analyses (OLS), were employed to disentangle the different contributors to visual response dynamics and estimate sources of visual response variance.

In a first approach, for each respective animal, the global signal and visual predictor were either separately or simultaneously regressed. The signal from visually activated areas (binary mask of significant group-level activations from GLM-based analysis) and the global signal across all brain areas were then calculated for all subjects. These time series and the QPP correlation vector were collected across all trials (n = 24 animals x 10 trials). The resultant distributions at each time point were analyzed and visualized as peri-event time traces, normalized to the 10s OFF period (Z-scoring procedure) prior to stimulation. Significant activations or de-activations at each time point were evaluated by one-sample T-tests (H0 distribution: μ = 0 and σ = sample σ) and FDR-corrected for the number of evaluated time points [n=90*s*/(0.5*s*/*TR*)] within each trial). Specifically, in these analyses, the purpose was not to directly compare the extent of statistical differences between various regression approaches, but rather to determine if there were significant increases or decreases in brain area time series and (particularly) QPP correlation vectors with regard to a zero-mean distribution. For visualisation, the QPP correlation vector was shown as mean Pearson correlation (ρ) rather than Z-score unit.

In a second approach, differences between visual response means were evaluated for stratified stimulation trials (all animals and trials), in different regression analyses. One-way analysis of variance (ANOVA) tests were performed for each time point during stimulation and model p-values were FDR corrected for the number of evaluated time points [n=30*s*/(0.5*s*/*TR*)]. Post-hoc tests were Bonferroni corrected.

## Acknowledgements

We thank Alessandro Gozzi, Daniel Barragan, Daniele Marinazzo and Xin Yu for their valuable feedback. This work was supported by the interdisciplinary PhD grant (ID) BOF DOCPRO 2014 (granted to M.V.) and further partially supported by funding received from: the European Union’s Seventh Framework Programme (FP7/2007-2013; INMiND) (grant agreement 278850, granted to AVdL.), the molecular Imaging of Brain Pathohysiology (BRAINPATH) and the Marie Curie Actions-Industry-Academia Partnerships and Pathways (IAPP) program (grant agreement 612360, granted to A.VdL.), Flagship ERA-NET (FLAG-ERA) FUSIMICE (grant agreement G.0D7615N), stichting Alzheimer onderzoek (SAO-FRA) (grant agreement 13026, granted to A.VdL.), the Flemish Impulse funding for heavy scientific equipment (granted to A.VdL.), the Fund for Scientific Research Flanders (FWO) (grant agreements G.048917N, G.057615N and G.067515N), the National Institutes of Health (NIH) (grant agreements R01MH111416-01 and R01NS078095), the National Science Foundation (NSF) (grant agreement BCS INSPIRE 1533260), and the ISMRM Research Exchange Program (granted to M.E.B.). The computational resources and services used in this work were provided by the HPC core facility CalcUA of the Universiteit Antwerpen, the VSC (Flemish Supercomputer Center), funded by the Hercules Foundation and the Flemish Government – department EWI.”

## Authors’ contributions

M.E.B. designed research, performed experiments, designed analysis, performed analysis, wrote paper. J.B. designed analysis and contributed valuable discussion. A.B., A.K, and W-J. P. contributed valuable discussions. R.H., V.V. and J.V.A. provided experimental and analytical support. A.V.D.L., S.D.K., and M.V. designed research and analysis. G.A.K. designed research, designed analysis and wrote paper. S.D.K., M.V and G.A.K. contributed equally to this work.

---

**Video 1.** QPP1 temporal evolution displayed per TR (0.5s) over a duration of 3s. Maps display Z-scores [n = 71 scans; Z-test with H0 through randomized image averaging (n=1000), FDR p<10^−7^, cluster-correction 4 voxels].

**Video 2.** QPP2 temporal evolution displayed per TR (0.5s) over a duration of 9s. Maps display Z-scores [n = 71 scans; Z-test with H0 through randomized image averaging (n=1000), FDR p<10^−7^, cluster-correction 4 voxels].

**Video 3.** QPP3 temporal evolution displayed per TR (0.5s) over a duration of 9s. Maps display Z-scores [n = 71 scans; Z-test with H0 through randomized image averaging (n=1000), FDR p<10^−7^, cluster-correction 4 voxels].

**Video 4.** Global signal temporal evolution displayed per TR (0.5s) over a duration of 9s (global signal peak occurs at 2s; image range [−2:7s] was chosen through cross-correlation of the global signal with QPP1-3 time series). Maps display Z-scores [n = 71 scans; Z-test with H0 through randomized image averaging (n=1000), FDR p<10^−7^, cluster-correction 4 voxels].

