## Supplementary for "Resting brain fluctuations are intrinsically coupled to visual response dynamics"

### Supplementary Material

#### Supplementary Methods

##### Animal handling in the scanner

Animals were anesthetized with 3.5% isoflurane and maintained at 2% during handling. Heads were fixed with bite and ear bars. Moldable earplugs were positioned on top of the ear bars in order to provide hearing protection. Ophthalmic ointment was applied to the eyes. Animal core body temperature was measured with a rectal temperature probe and kept stable at 37 °C via hot air supply (MR-compatible Small Animal Heating System, SA Instruments, Inc.). A pressure sensitive pad was used to assess breathing rate, and a fiber-optic pulse oximeter placed over the tail, to assess heart rate and $O_2$ saturation (MR-compatible Small Animal Monitoring and Gating system, SA Instruments, Inc.).

##### MRI procedures and registration

Rapid Acquisition with Refocused echoes (RARE) images were acquired in three orthogonal directions (repetition time (TR) 2500ms, effective echo time (TE) 33ms, 16 slices of 0.4 mm), to enable reproducible slice positioning. B0 field maps were acquired, followed by local shimming. Functional MRI scans were acquired with a gradient-echo echo-planar imaging (EPI) sequence (field of view (27x21x6) $mm^3$, matrix dimensions [90x70x12], slices thickness 0.4mm, slice intersperse 0.1mm, flip angle 55°, bandwidth 250kHz, TR 500ms, TE 14ms). In each scan session, a T2-weighted 3D anatomical scan was acquired (RARE, TR 1800ms, TE 6ms, RARE factor 16, spatial resolution (0.078 x 0.078 x 0.31) $mm^3$). The open source registration toolkit ANTs was used to construct a study-based 3D T2 RARE template. The 2D EPI-based study template was then registered to the Allen brain atlas (1) with ANTs using a two-stage registration procedure. First, registration of the masked and debiased 3D study template to the atlas was performed, and second, registration of the masked and debiased 2D EPI study template to the 3D study T2 RARE template was performed. Using the 2-stage registration parameters, EPI template space and related significance maps were forward-transformed into atlas space.

##### Frame-wise displacement

Frame-wise displacement (FD), for all subject in the LA short TR data set, was calculated at each point by taking the sum of absolute backwards looking temporal derivatives for all three motion time series (2). To compute rotational displacement and convert degrees to millimeters, an assumption was made where the mouse brain is considered as a sphere with a diameter of 10mm.

**Resting state independent component analysis**

To obtain intrinsic connectivity networks (ICN), group ICA was performed on all rsfMRI scans (n = 71; **Supplementary Table 1**) using the GIFT toolbox (v4.0b) (Calhoun et al., 2004). The number of independent components was set to N=5. If the number of components was increased beyond five, anti-correlation diminished and the DMN became split into an anterior and posterior component. The ICA was run on variance-normalized data (z-scoring procedure), filtered between 0.008-0.2Hz, using the Infomax algorithm with no auto-filling of data reduction values. Stability analysis was performed using the ICASSO algorithm, rerunning the ICA 20 times with a minimal cluster size of 10 and maximal of 20. Using the built-in GIFT functionality, SPM12 was used to obtain T-contrast images per subject for all components. Second level one sample T-tests were performed for all components to obtain significant group level ICNs [$p<10^{-5}$, FDR, cluster-correction 4 voxels).

**Quasi-periodic pattern selection – determining window size**

For each set of QPPs, at a respective window size, QPPs were first grouped into opposing phase observations. Opposing phase implies that within a set, QPPs displayed similar involvement of spatial areas, but with inverted timing of activation and deactivation. This heuristic assumption was inspired by prior observations (3, 4), and overall visual inspection of the current findings. In order to phase QPPs, the signal of a specific set of brain areas over the duration of the QPP was used. Specifically, DMN-like areas have been consistently observed as a component of QPPs. Therefore, a mask of the DMN-like ICN was derived from the ICA analysis (**Supplementary Figure 1**) and was used to average the intensity of all DMN voxels at each time frame within the QPPs. Using these time courses, QPPs were phase sorted according to a previously described strategy [cfr. (3)]. From the phase sorted groups at each window size, a single QPP was obtained by selecting the one that displayed the highest value of summed STC peaks [cfr. (5)]. As QPPs at smaller window sizes are expected to be parts of longer bi-phasic QPPs, a previously described algorithm was used to select the most representative window length for the identified QPPs (3). This algorithm identifies the amount of overlap between QPPs of different lengths, termed the fractional average correlation (FA), by calculating image correlation at different temporal lags. Then, it identifies the window length after which longer variants of the same QPP no longer substantially increase the FA value. This determines the most representative window length and allows selection of QPPs for further investigation. For the current study, this cut-off point was 9s.

**Power spectral analysis**

Subject-specific power spectra, for any respective time series, were obtained via multitaper power spectral density estimation, using the full spectral range (Nyquist frequency = 1Hz). Average power spectra were determined for rsfMRI and visual-evoked fMRI scans receptively. On the average spectra, power-law distributions were fit via ordinary least squares (OLS) in the 0.01-0.2Hz range. The fit was determined via the expression P = $1/f^{\beta}$, where P is power, f is frequency, and $\beta$ is the power-law exponent that informs on the slope of the fit (P and f are log-scaled). The power law function estimates scale invariance of the system under consideration and can be used to evaluate a lack of periodicity in an fMRI-derived power spectrum (6). Deviation of the power spectrum from the fit indicates signal (quasi-)periodicity.

**Haemodynamic response function**

Currently, no true mouse HRF is available in the literature, but prior work has indicated fast and sharp neurovascular coupling in the mouse brain (7–9). Particularly, the observations made by Lebhardt and colleagues (2015) were here parameterized by a mixture of two gamma variate functions within SPM, using the function “spm_hrf” (settings: response delay = 2s, undershoot delay = 6s, response dispersion = 0.5, undershoot dispersion = 1, response/undershoot ratio = 12, onset = 0s, kernel length = 6s).

#### Supplementary Text

**Supplementary Results.** Intrinsic brain response to visual stimulation

After regression of the visual signal predictor, a 9s STP was observed in the visual-evoked fMRI scans [in addition to the 3s STP; (**Figure 3B**)] that appeared similar to QPP3 identified from resting state scans (**Supplementary Figure S4**). This particular pattern displayed a trial-locked behaviour (**Supplementary Figure S6**), but this was less pronounced compared to that observed for the 3s STP (**Figure 3B**). The response dynamics of the 9s STP in the visual fMRI scans could be accounted for by regression of the short 3s STP (**Supplementary Figure S6**). No significant trial correlation could be determined for QPP2 (**Supplementary Figure S6**). Overall, these findings suggest that only QPP1 appeared to be clearly evoked through visual stimulation.

**Supplementary Discussion. Neuromodulation and the reticular activating system**

Activity in the reticular formation and neuromodulatory brain regions was observed in several of the presented findings. The reticular formation, and its regulation of the ascending reticular activating system (ARAS), is responsible for promoting wakefulness and attention, mediated via either a thalamic pathway that further projects to cortical areas, direct projections to the cortex, or a hypothalamic / basal cholinergic forebrain pathway that projects to cortical areas. The ARAS is marked by its inclusion of a range of neuromodulatory nuclei such as raphe nucleus (serotonergic), locus coeruleus (adrenergic), nucleus basalis (cholinergic), the penduncHulopontine nucleus (cholinergic), and ventral tegmental area (dopaminergic). The latter are reciprocally interconnected and may affect each other via different pathways, yet, may serve separate functions and innervate specific brain areas. Overall, there is widespread involvement of different modulatory systems in the ARAS [for reviews on the subject matter, please refer to (10–14)]. It has been suggested that these structures are natural rhythm generators that may provide infraslow patterned input to the brain (15). At the same time, they allow selective adaptation to modulate brain state and direct attention to sensory stimuli (16–20). The observation of rhythmic activations and deactivations in the reticular formation might be understood from this structure's role in regulating (de-)synchronized and REM sleep. Different regions of the reticular formation may display fluctuations in tonic inhibition and activation that in concert can be transformed into rhythmic activity. Cholinergic reticular neurons, vestibular nucleus neurons and vestibulo-oculomotor neurons may for instance generate bursts of REM, which project activity to ascending and descending motoric systems (21). Similarly, pontogeniculooccipital (PGO) waves may originate in a burst from the pons, travelling to geniculate and occipital visual cortical areas, and play an important role in the regulation of eye saccades (22). Interestingly, it has been suggested that PGO waves may also affect other brain areas, particularly hippocampus, entorhinal cortex and Piriform cortex (23). In this review, Gott et all. (2017) point out that PGO waves may display even more widespread cortical involvement and hint that its functional role may relate to the DMN and the internal simulation of external sensory information. The DMN, and its anti-correlated network, observed in the current study are reflecting such descriptions (**Figure S1**). It further stands out that visual input may directly affect activity in the reticular formation and promote arousal (24). Of particular relevance, it has been suggested that PGO waves reflect the startle network of the brain (25). The reticular formation also converts external stimuli into input to the locus coeruleus, which in turn may facilitate sensory processing, by resetting cortical activity and adapting the brain state for a behavioural response (26, 27).

**Supplementary Discussion. Resting state networks**

Resting state networks displayed close similarity to preceding mouse literature (28–30), revealing the DMN, lateral cortical network, (hypothesized to reflect the TPN), somatomotor network, sensory networks, forebrain, and cerebellum. Interesting additional features were observed, which may derive from the unique nature of the large-sample whole brain rsfMRI acquisitions at a temporal resolution of 0.5s. Overall, networks displayed a high degree of large-scale anti-correlation, even though no global signal regression was performed. The DMN-like network displayed anti-correlation with bulbus, ventral striatum and ventral hippocampus. These areas are part of an anatomical striatal-limbic network in rodents (31–33). Functional coupling between hippocampus and the bulbus has also been linked to respiration-driven rhythmicity in awake rats (34). The DMN further appeared anti-correlated with a focal region in the reticular formation, while other parts of the reticular formation were correlated. A study in humans indicated that a seed-based FC map indicating the DMN was obtained when a seed was placed in the reticular formation, which further appeared to be a causal driver of DMN activation (35). Interpretations for the deactivation in the reticular formation were presented above. The TPN-like network displayed anti-correlation with brain areas that comprise an anterior dorsal part of the DMN. Similar observations have been made in mice under similar anesthesia (36). Also in humans, it has been indicated that the DMN is comprised of several sub-networks (37–39), and that the dorsal section of the DMN naturally displays more anti-correlation with the TPN when no global signal regression is performed (40). Other anti-correlated areas with the TPN were the medial bulubus, Pallidum, septal nuclei, and reticular formation. Deactivation of the cholinergic basal forebrain (pallidum) may be understood in light of a recent study which showed that a specific pathway of this brain area couples local gamma band activity to gamma band activation in the DMN during resting conditions (41). Deactivation in the reticular formation, similar as was observed for the DMN, might be understood as combination of different pathways (e.g. added inclusion of cholinergic basal forebrain) that can differentially modulate variable networks. Septal nuclei are important for regulating hippocampal activity, a function that may be mediated through serotonergic inputs to the septal nuclei and in turn cholinergic modulation of the hippocampus (42). Given that the hippocampus is part of the DMN, this observation might indicate a deactivation of posterior DMN components. ICN 3 displayed a somatomotor network, which was anti-correlated with visual cortex, motor cortex, Entorhinal/Parahippocampal cortex, medial Pallidum and Septal nuclei. The listed cortical areas are anatomically connected and form a functional network that serves visuo-spatial processing and episodic memory encoding (43). A role for pallium and septal nuclei was provided above. ICN 4 displayed a sensory network composed of olfactory systems, auditory cortex and visual cortex, correlated with the hippocampus and entorhinal cortex, and anti-correlated with the brainstem and midbrain. Low frequency optogenetic stimulation of the ventral posteriomedial midbrain has been shown to cause widespread BOLD activation in sensory brain areas in rats (44). Low frequency optogenetic stimulation of the hippocampus in turn has been shown to cause widespread BOLD activations in cortical areas, particularly so visual cortices (45). ICN 5 displayed a clear cerebellar component (28), which was anti-correlated with a large-scale forebrain system, mainly comprising striatal areas, prelimbic areas and prefrontal cortex. Overall the observed ICNs appear to capture sensible functional BOLD networks that provide new insights on top of existing mouse literature.

#### Supplementary tables and figures

| | Study Design | | | |
|---|---|---|---|---|
| | Session 1 (week 0) | | Session 2 (week 2) | |
| | 30-40m | 40-55m | 30-40m | 40-55m |
| subj001-012 | rsfMRI 1 | rsfMRI 3 | rsfMRI 4 | visual fMRI |
| subj013-024 | rsfMRI 2 | visual fMRI | rsfMRI 5 | rsfMRI 6 |

**Table S1. Overview of experimental design.** Matching colours mark consistent scan types.

| | mean framewise displacement (mm) | | | |
|---|---|---|---|---|
| | Session 1 (week 0) | | Session 2 (week 2) | |
| | rsfMRI | rsfMRI | rsfMRI | visual fMRI |
| subj001 | 0.0425 | 0.0442 | 0.0425 | 0.0443 |
| subj002 | 0.0415 | 0.0415 | 0.0393 | 0.0416 |
| subj003 | 0.0452 | 0.0429 | scan error | 0.0416 |
| subj004 | 0.0421 | 0.0406 | 0.0511 | 0.0484 |
| subj005 | 0.0381 | 0.0365 | 0.0344 | 0.0361 |
| subj006 | 0.0381 | 0.0386 | 0.0428 | 0.0437 |
| subj007 | 0.0422 | 0.0409 | 0.0358 | 0.0377 |
| subj008 | 0.0457 | 0.0437 | 0.0361 | 0.0378 |
| subj009 | 0.0433 | 0.0452 | 0.0366 | 0.0367 |
| subj010 | 0.0472 | 0.0449 | 0.0337 | 0.0350 |
| subj011 | 0.0393 | 0.0397 | 0.0388 | 0.0392 |
| subj012 | 0.0432 | 0.0432 | 0.0364 | 0.0366 |
| | rsfMRI | visual fMRI | rsfMRI | rsfMRI |
| subj013 | 0.0488 | 0.0490 | 0.0376 | 0.0386 |
| subj014 | 0.0564 | 0.0583 | 0.0383 | 0.0419 |
| subj015 | 0.0399 | 0.0370 | 0.0345 | 0.0355 |
| subj016 | 0.0378 | 0.0374 | 0.0445 | 0.0425 |
| subj017 | 0.0413 | 0.0397 | 0.0348 | 0.0354 |
| subj018 | 0.0378 | 0.0393 | 0.0388 | 0.0402 |
| subj019 | 0.0355 | 0.0357 | 0.0357 | 0.0385 |
| subj020 | 0.0406 | 0.0397 | 0.0336 | 0.0340 |
| subj021 | 0.0380 | 0.0395 | 0.0357 | 0.0370 |
| subj022 | 0.0408 | 0.0418 | 0.0358 | 0.0368 |
| subj023 | 0.0383 | 0.0402 | 0.0375 | 0.0390 |
| subj024 | 0.0388 | 0.0389 | 0.0441 | 0.0367 |

**Table S2. Mean frame-wise displacement.** Colour codes in Table S1. For one animal, a scan reconstruction error occurred (red text).

| Group comparisons |
| --- |
| session 1 rsfMR 1+2 versus session 2 rsfMRI 4+5<br>test: voxel-wise paired T-test of RSNs<br>correction: FDR p < 0.05<br>outcome: no differences |
| session 1 rsfMR 1 versus session 1 rsfMRI 3<br>test: voxel-wise paired T-test of RSNs<br>correction: FDR p < 0.05<br>outcome: no differences |
| session 2 rsfMR 5 versus session 2 rsfMRI 6<br>test: voxel-wise paired T-test of RSNs<br>correction: FDR p < 0.05<br>outcome: no differences |
| session 1 visual fMR versus session 2 visual fMRI<br>test: voxel-wise two-sample T-test of visual activations<br>correction: FDR p < 0.05<br>outcome: no differences |

**Table S3. Group comparison.** Each compartment indicates tests that compared similar scan types. These were evaluated to determine if data pooling was appropriate. No images are shown, because no significant differences were apparent on networks or activation maps.

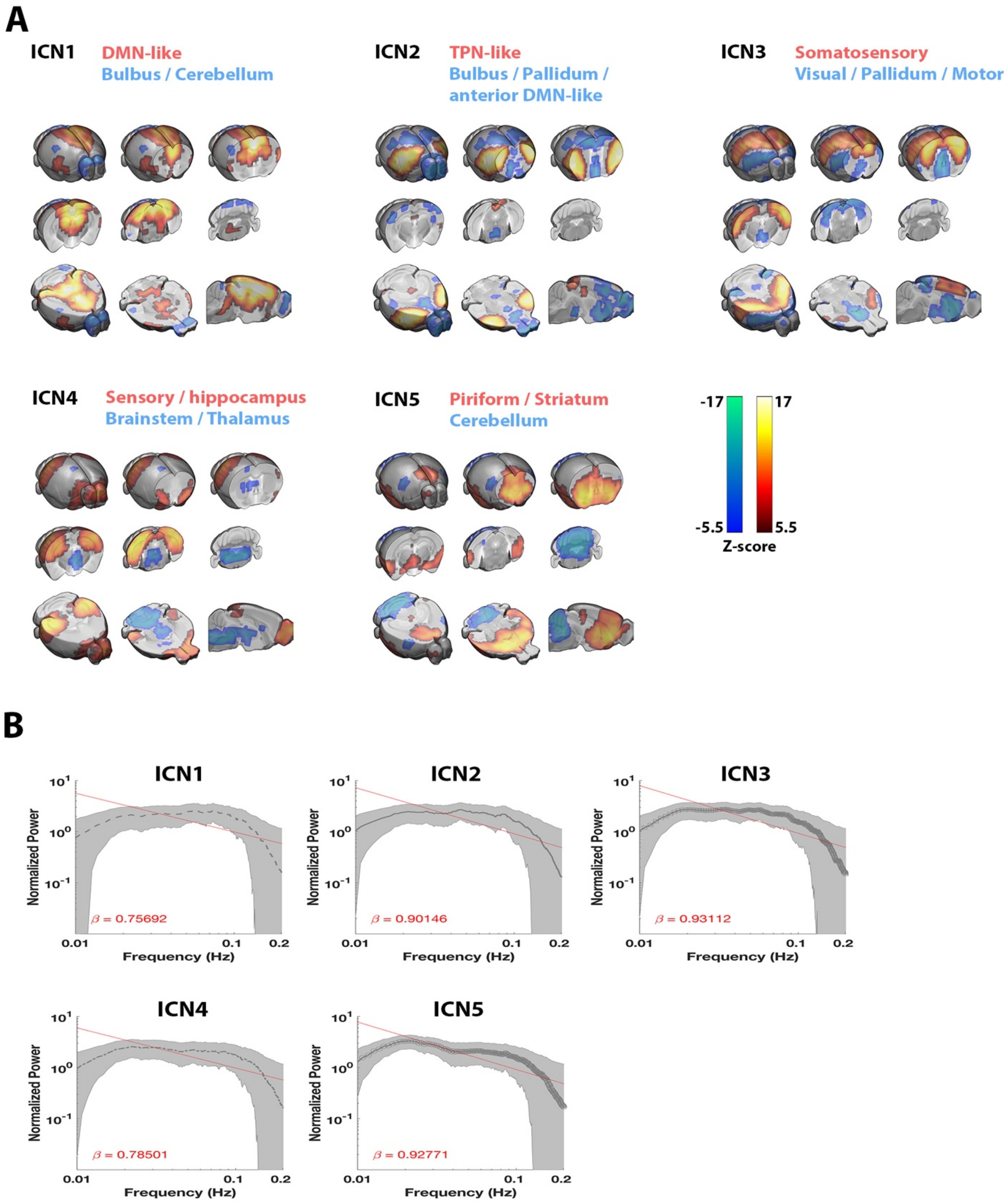


**Figure S1. Intrinsic connectivity networks. A)** Five major intrinsic connectivity networks (ICN) were observed during the resting state, displaying wide-spread anti-correlation. n = 71 scans. Maps display Z-scores (first level GLM; second level one sample T-test; T-scores normalized to Z-scores; FDR $p<10^{-7}$). **B)** Log-log power spectra for each ICN. A Power-law distribution was fit via OLS (red line), and the slope (power law exponent $\beta$) is shown. Traces display mean, patches display STD.

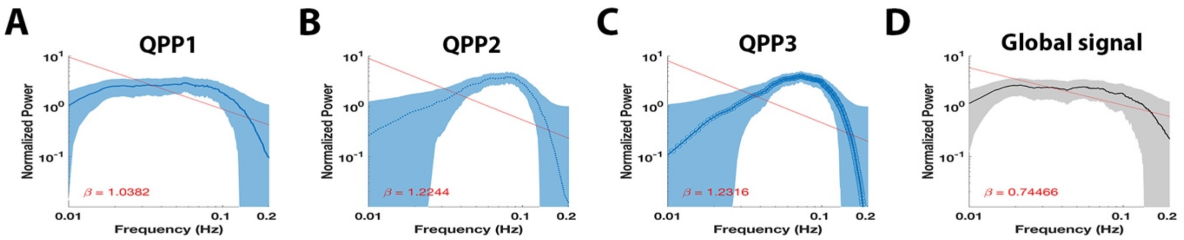


**Figure S2. QPP and global signal power spectra**

All three QPPs displayed quasi-periodic temporal characteristics, which were less pronounced in QPP1 **(A-C)**. Further, the global signal displayed a power spectrum similar as observed for QPP1, but with a further reduced quasi-periodic component, approaching aperiodicity **(D)**. **A-D)** Log-log power spectra. A Power-law distribution was fit via OLS (red line), and the slope (power law exponent β) is shown. Traces display mean; Patches display STD.

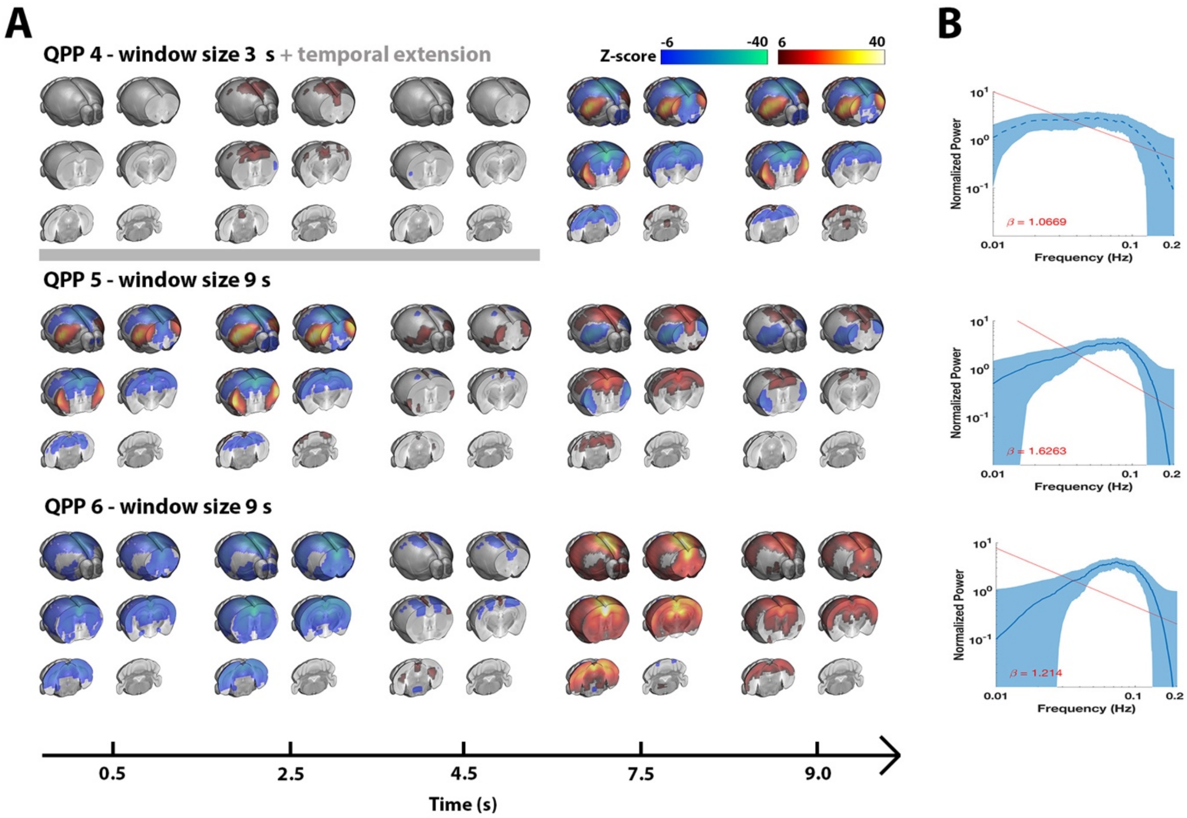


**Figure S3. Opposite phase QPPs.** This figure complements Figure 1 in the main text and shows the opposite phase variants of QPP1-3. **A)** QPPs are displayed on the same time axis (alignment through cross-correlation of QPP correlation vectors). Additional images before the 3s core of QPP4 were averaged to visualise likelihood of preceding activations. Maps display Z-scores [Z-test with H0 through randomized image averaging (n=1000), FDR $p<10^{-7}$]. **B)** Log-log power spectra. A Power-law distribution was fit via OLS (red line), and the slope (power law exponent $\beta$) is shown. Blue traces display mean, blue patches display STD.

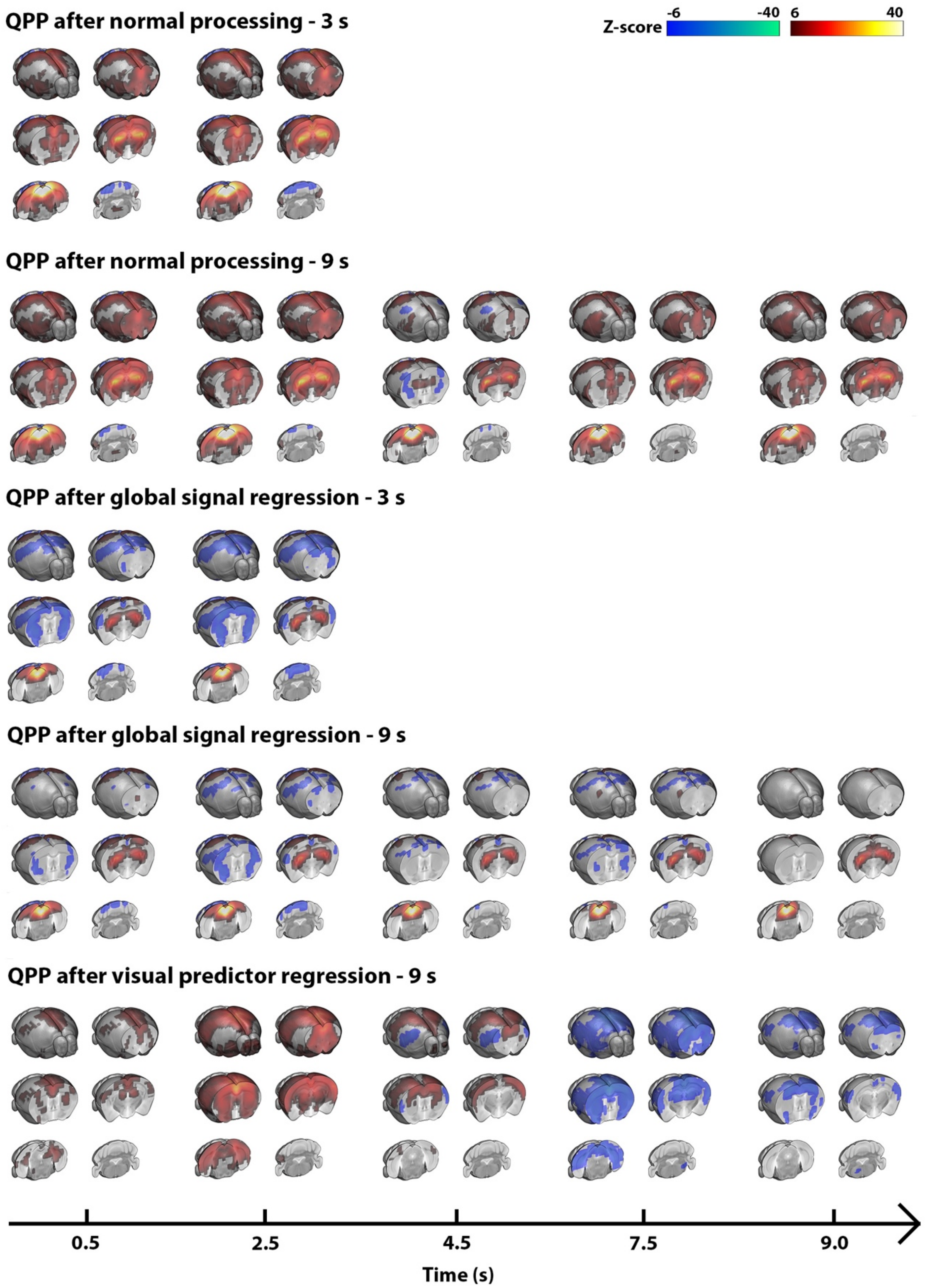


**Figure S4. QPPs during visual stimulation.** This figure complements Figure 3 in the main text and shows additional QPPs that could be determined from visual fMRI scans, using various regression methods (indicated atop respective images). The lowest shown 9s QPP, obtained after visual predictor regression, was similar to QPP3 in the resting state. N = 24 scans. Maps display Z-scores [Z-test with H0 through randomized image averaging (n=1000), FDR $p<10^{-5}$].

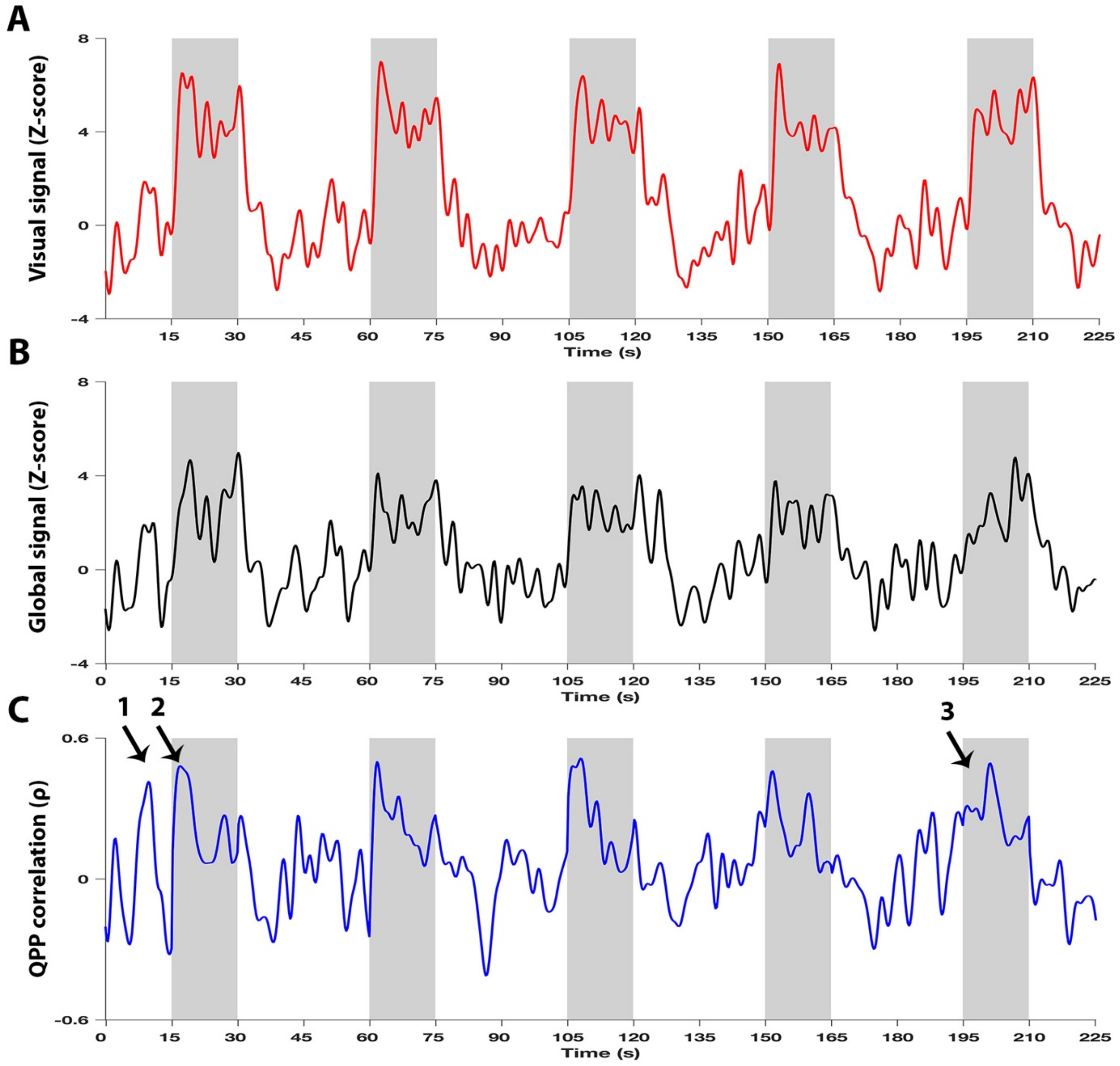


**Figure S5. Single subject time courses during visual-evoked fMRI.** A 7-minute excerpt is shown from the visual-stimulation fMRI recordings in a single mouse (5 trials). Red trace shows signal from visual areas, black the global signal, blue the QPP correlation vector. Black arrows indicate noteworthy observations: 1) strong QPP correlation peak during the period prior to a stimulation (OFF); 2) Strong QPP correlation peak at the start of stimulation (ON); 3) No QPP correlation peak at the start of stimulation (ON).

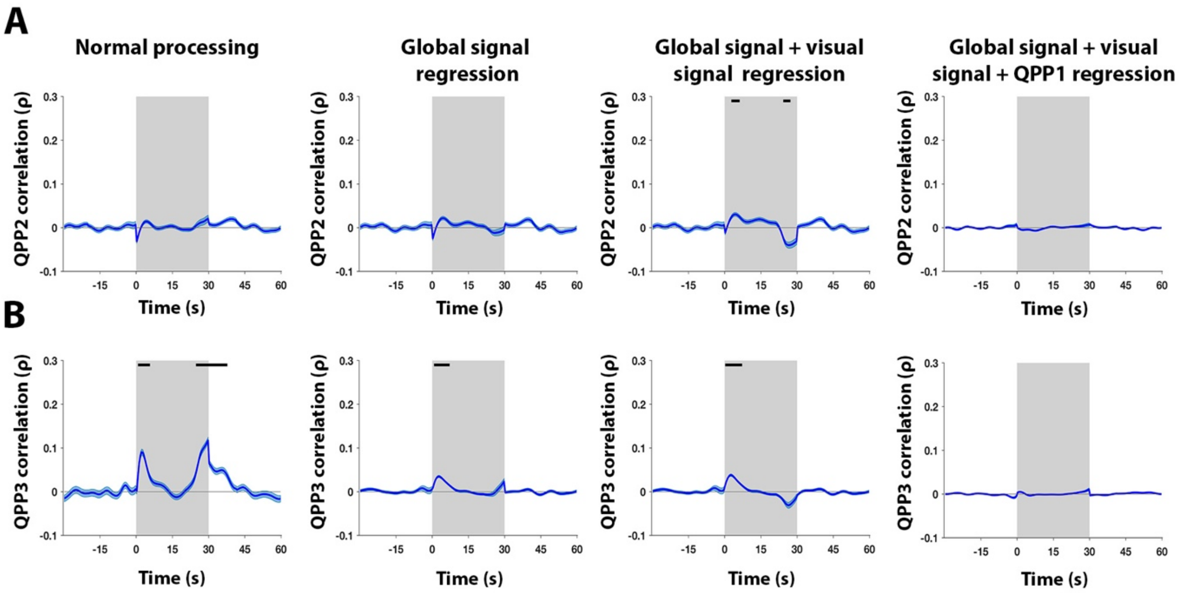


**Figure S6. Trial-locked behaviour of QPP2 and QPP3.** This figure complements Figure 3 in the main text and shows trial-average behaviour of QPP2 and QPP3. Here, QPP2 is the pattern observed during resting state, correlated with the task fMRI image series (it was not directly observed in task fMRI scans). QPP3 is the STP directly determined from the task fMRI image series, which displayed high similarity to resting state QPP3. n = 24 animals x 10 trials. Grey areas mark trials (ON periods), traces show mean (BOLD time courses demeaned and variance normalized to 10s OFF period prior to stimulation), patches show STD, black bars mark significance (one sample T-test, FDR $p<10^{-5}$). Note the lack of correlation for QPP2 and the weaker correlation of QPP3 in contrast to the 3s STP indicated in Figure 3. Further, note the lack of correlation for both QPP2 and QPP3 when QPP1 is regressed from the respective image series.

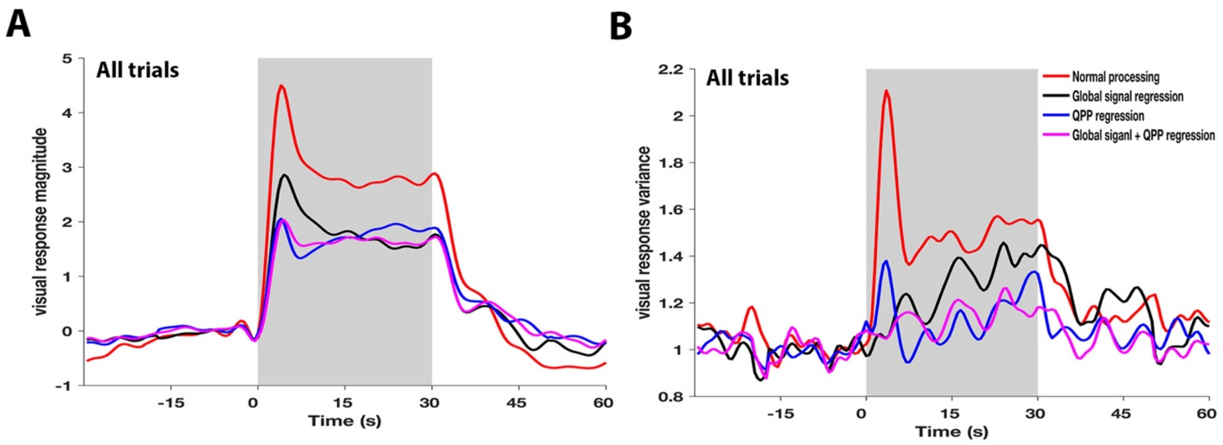


**Figure S7. Visual response magnitude and variance under different conditions of intrinsic signal regression.** Across all trials, QPP or global signal regression substantially lowered visual response amplitudes **(A)**. After QPP regression, initial peak responses amplitudes were almost completely removed. These regressions also significantly lowered visual response variance **(B)**. Independently, QPP or global signal regression, significantly lowered variance during visual stimulation, but combined QPP and global signal regression even further reduced variance so that almost no more differences were apparent with rest blocks. **A-B)** n = 24 animals x 10 trials. Time traces are demeaned and variance normalized to 10s OFF period prior to stimulation.
